# Spatial proteomic analysis of a lung cancer model reveals regulatory T cells attenuate KRAS-G12C inhibitor-induced immune responses

**DOI:** 10.1101/2024.04.11.588725

**Authors:** Megan Cole, Panayiotis Anastasiou, Claudia Lee, Chris Moore, Edurne Mugarza, Martin Jones, Karishma Valand, Sareena Rana, Emma Colliver, Mihaela Angelova, Katey S.S. Enfield, Alastair Magness, Asher Mullokandov, Gavin Kelly, Tanja D. de Gruijl, Miriam Molina-Arcas, Charles Swanton, Julian Downward, Febe van Maldegem

## Abstract

We recently showed that lung tumor specific KRAS-G12C inhibition causes remodelling of the tumor immune microenvironment from cold to hot. As a result, KRAS-G12C inhibition is able to synergise with anti-PD-1 treatment, but only in tumor models that were already moderately responsive to immune checkpoint blockade at baseline. To investigate mechanisms that restrain immunotherapy sensitivity in non-responsive tumors, we used multiplex imaging mass cytometry to explore spatial patterns in the tumor microenvironment of the highly immune evasive KRAS mutant murine Lewis Lung Cancer model. Clustering of close neighbour information per cell allowed characterisation of spatial patterns or ‘communities’ in the tissue. We identified a community harbouring features of localised T-cell activation, where CD4^+^ and CD8^+^ T cells and dendritic cells were gathered together. KRAS-G12C inhibition led to increased expression of PD-1 on T cells, CXCL9 expression by dendritic cells, together with increased proliferation and potential cytotoxicity of CD8^+^ T cells, indicating an effector response. However, we also observed a high incidence of regulatory T cells (Tregs) within this community, which had frequent contact with effector T cells, suggesting that Tregs may be able to dampen anti-tumoral immune responses following KRAS-G12C inhibition. Similar communities were detected in human lung adenocarcinoma clinical samples. Depleting Tregs *in vivo* with anti-CTLA-4 antibody rescued the anti-tumor immune response and led to enhanced tumor control in combination with anti-PD-1 and KRAS-G12C inhibitor. We therefore propose use of KRAS-G12C inhibitor in combination with Treg depletion as a therapeutic opportunity that increases anti-tumoral immune responses and initiates tumor regression.

**One sentence summary:** Spatial analysis identified regulatory T cells as potential source of local T cell repression, mediating resistance to KRAS-G12Ci and anti-PD1 therapy.

## INTRODUCTION

Recent years have seen a transformation in the treatment of non-small cell lung cancer (NSCLC), with the introduction of immune checkpoint blockade, which has increased survival rates of patients with a previously poor prognosis. Despite this, only a subset of patients responds and many responders acquire resistance over time (*1, 2*). In 2021 a further breakthrough occurred when the RAS inhibitor sotorasib was approved for the treatment of locally advanced or metastatic KRAS-G12C mutant NSCLC. This followed successful clinical trials where 80% of patients achieved temporary disease control following sotorasib treatment. However, despite a modest improvement in progression free survival, sotorasib gave no improvement in overall survival compared to docetaxel (*3*), demonstrating its limitations for use as a monotherapy. Therefore, strategies for combination with other therapies are being urgently sought (*4, 5*).

The importance of the immune system in the response to KRAS-G12C inhibition was revealed when Canon *et al.* showed that T cell presence was essential for durable responses in subcutaneous tumors of the colon cancer model CT26 treated with sotorasib (*6*). Additionally, Briere *et al.* using the same model demonstrated a switch in the tumor microenvironment (TME) from immunosuppressive in the vehicle setting, with high presence of M2 macrophages and MDSCs, to favouring anti-tumoral immune response following KRAS-G12C inhibition with adagrasib, another clinically approved KRAS targeted drug (*7*). CT26 tumors show an immune hot TME (i.e. high T cell infiltration rates) and are responsive to single agent immune checkpoint inhibition (ICI). Our previous work also revealed remodelling of the TME following KRAS-G12C inhibition in multiple lung tumor models, including the immune cold orthotopic Lewis Lung (3LL) murine NSCLC tumor model (*8, 9*).

KRAS-G12C inhibitors were also shown to synergise well with ICI in lung cancer models, but only in immune hot TME settings (*8, 10*). These findings highlight that while KRAS-G12C inhibitors specifically target tumor cells, this results in profound secondary effects on the TME, and T cells are crucial for durable responses.

Our previous analysis established that following seven consecutive days of treatment with the KRAS-G12C inhibitor MRTX1257, although 3LL tumor growth was inhibited, tumors did not regress, indicating that KRAS-G12C inhibitors alone are not sufficient to cause tumor regression in this model (*9*). Combination of this KRAS-G12C inhibitor with anti-PD-1 therapy did not increase responses in this model (*8*). This is reflective of the failure to see a beneficial effect on response of combined KRAS inhibition and PD-1 blockade in the clinical setting, either due to combination toxicities or lack of efficacy (*11*). The unmet need for new combinatorial treatment options which improve T-cell mediated anti-tumoral immune responses in immune cold tumor models has hence become increasingly clear. We therefore used the 3LL immune evasive orthotopic lung tumor model, in which effector immune cells are excluded from the tumor, to seek more effective therapeutic combinations with KRAS-G12C inhibition.

We used imaging mass cytometry (IMC) to analyse the make-up of these tumors in situ. This method is particularly useful for study of the TME due to its ability to capture up to 40 markers simultaneously. Spatial information remains intact, meaning that cell phenotypes can be analysed in the context of their spatial neighbours (*12*). Obtaining spatial information is essential when studying the TME, as it provides insight into the cellular interactions dictating local immune activation or suppression, as mechanisms of immune response or resistance to treatment, and has been shown to improve prediction of clinical outcomes compared to cell frequency alone in NSCLC patients (*13*).

Here we present data on the identification of neighbourhood communities through single cell spatial analysis to investigate which cellular interaction patterns may restrain anti-tumoral immune responses in the 3LL tumor model. A community resembling a T cell activation hub was identified, where regulatory T cell interactions may play a key role in dampening anti-tumoral immune responses following KRAS-G12C inhibition. In parallel, cellular community analysis of treatment-naïve human lung adenocarcinoma patient samples from the RUBICON TRACERx cohort, suggested that similar local Treg control may be restraining immune responses in a subset of patients.

Altogether, this led us to explore the effect of combining KRAS-G12C inhibitor with a Treg depleting anti-CTLA-4 antibody in the in vivo orthotopic setting, where markedly improved responses were noted. This opens up the perspective of combining KRAS-G12C inhibitors with Treg targeting to improve durable response rates.

## RESULTS

### Introduction to spatial communities and validation

Our previous IMC analyses on 3LL lung tumors have shown that there are clear patterns in the arrangements of cells in the lung cancer tissues, and changes to those patterns occur in response to KRAS-G12C inhibition (*9*). For example, we saw one subset of macrophages lining the tumor-normal interface, while another type of macrophages was intermixed with the tumor cells. Effector cells such as T cells and B cells were excluded from the tumor domain, while treatment with KRAS-G12C inhibitor MRTX1257 induced movement of T cells and antigen presenting cells into the tumor domain. We also noted that the phenotypes of cells differed depending on their location in the tissue and hypothesised that the local neighbourhood is likely to strongly influence the activation or inhibition of immune cells. Therefore, we adopted a cellular community analysis, to cluster cells based on the composition of their local neighbourhood (adapted with modifications from (*14*)). Analysis focused on two previously published datasets (*8, 9*), both generated from an in vivo experiment in which the Lewis Lung (3LL) carcinoma model was treated with the KRAS-G12C inhibitor MRTX1257 or Vehicle for 7 days prior to harvesting of the lungs. The tumors were stained with two partially overlapping antibody panels to give two datasets, with the dataset 2 panel being more T-cell oriented (Supp. Fig. 1a, b). The cell typing was derived from our previous analyses and based on lineage markers only, but independent of maturation or activation markers. Of note, the subsequent cellular communities were therefore blind to cell phenotypes.

We identified neighbours by a 15-pixel expansion of the cell boundary through segmentation in CellProfiler. A 15-pixel radius was chosen as it depicts the average size of a cell, and therefore cells identified as neighbours would be those ‘up to one cell away’. As a result, neighbourhoods were not equal in cell number, but instead reflected the local density surrounding each cell. Louvain clustering using Rphenograph was then run on the neighbour proportion information per cell to identify recurring spatial patterns in the tissue, labelled ‘spatial communities’ or just ‘communities’ in short (Fig. 1a). Graph building with a k-nearest neighbour input value of 250 yielded 62 communities for dataset 1, these were agglomerated to 30, and subsequently 18 communities. Agglomeration merged the communities with the lowest cell diversity, representing mainly small variations in the tumor cell neighbours (agglomerated community 3), while the highly diverse communities, such as those with a high proportion of immune cells, remained stable (Fig. 1b, Supp. Fig. 1c). We were most interested in the immune-dominated communities for this analysis and therefore decided 18 communities was optimal to carry forward for our investigation into anti-tumoral immune response.

**Figure 1.**
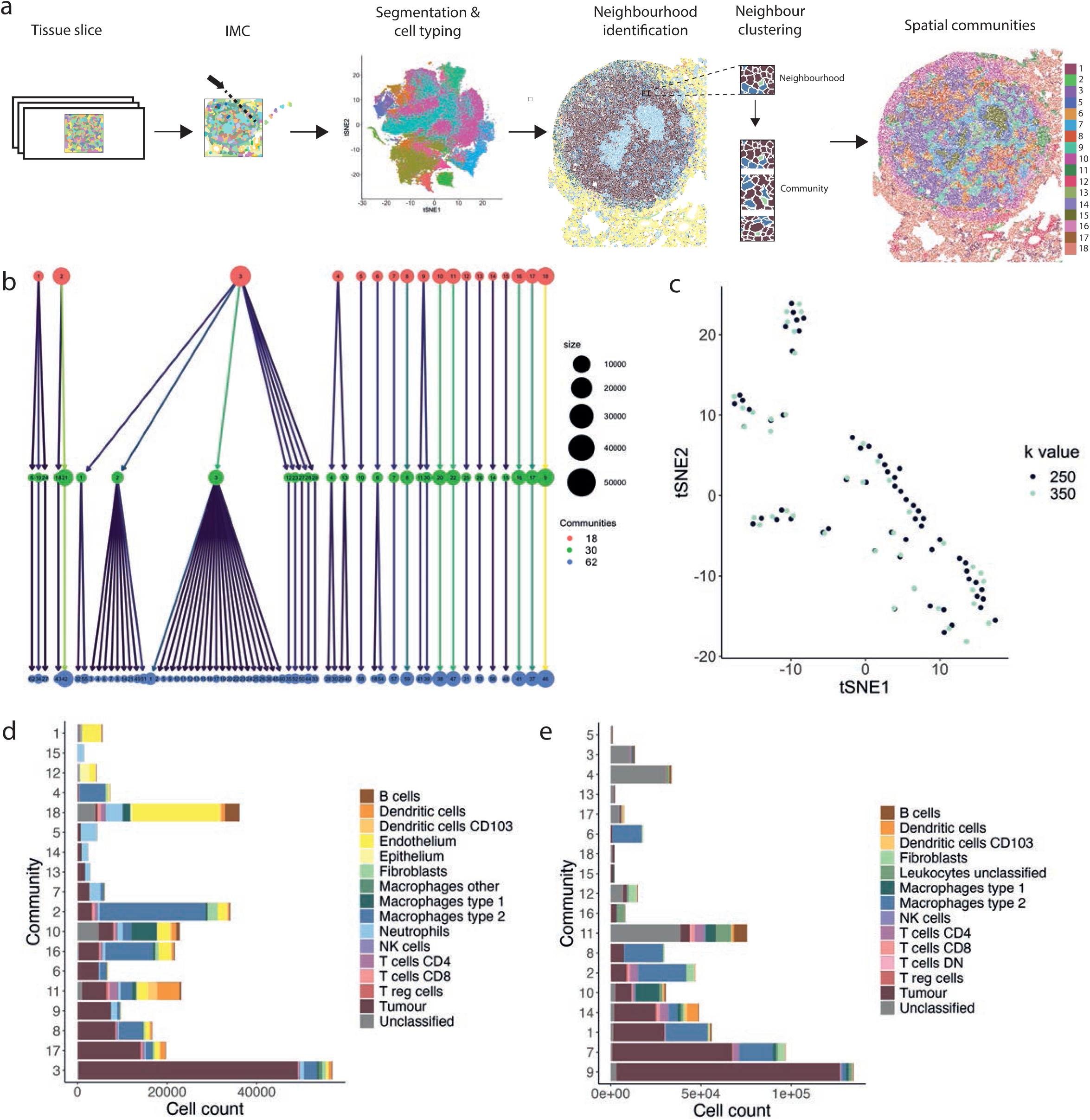
Introduction to spatial communities and validation. **a)** Workflow of generating spatial communities using the data generated from lung tissue slices. **b)** Cluster tree of 62 communities generated using Rphenograph with a k-input value of 250, agglomerated to 30 communities and subsequently agglomerated again to 18 communities, where each circle represents a community and lines indicate communities that were merged during agglomeration. **c)** tSNE plot of 62 communities generated with a k-input value of 250 and 47 communities generated with a k-input of 350 into Rphenograph using dataset 1, were tSNE analysis was run based on the proportion of each cell type contributing to each community. **d)** 18 spatial communities generated from clustering based on neighbour proportions of each cell type for **d)** dataset 1 and **e)** dataset 2, with the size of each bar representing cell count of that community and colours indicating the contribution of each cell type. Bars ordered by decreasing tumor cell count.

To determine whether our method to identify spatial communities was robust to altered clustering input conditions, we also ran Rphenograph clustering on the neighbourhood information for all cells in dataset 1 with a k-input value of 350. Dimensionality reduction using tSNE (t-distributed Stochastic Neighbour Embedding) of the 62 communities identified from k-input of 250 and the 47 communities identified from a k-input of 350 revealed very similar patterns of community phenotypes (Fig. 1c). Additionally, there were multiple overlaps between communities identified using the different k-values for clustering, suggesting matching phenotypes.

The 254 communities identified from neighbour clustering of dataset 2 using a k-input of 250 were also agglomerated to 18 communities to enable parallel analysis with the communities identified in dataset 1. The communities in both datasets varied largely in size, with some containing fewer than 2,000 cells, and others comprising over 50,000 cells (Fig. 1d, e). There was also differing heterogeneity of these spatial groups, with some being dominated by a single cell type, such as tumor cells in community 3 from dataset 1 and community 9 from dataset 2, whilst others comprised a mixture of various cell types in more balanced proportions (Supp. Fig. 1d, e).

Datasets 1 and 2 were based on different antibody panels and therefore not identifying all the same cell types. For example, lack of markers EPCAM and PECAM for dataset 2 meant endothelial and epithelial cells could not be identified and thus a large number of cells were labelled as ‘unclassified’. Nevertheless, a separate community clustering analysis on both datasets based on shared cell types only, demonstrated that the method to generate communities was stable to altered input data (Supp. Fig. 1f).

### Spatial communities reflect refined tissue architecture and have functional relevance

As the spatial communities are based on local neighbourhoods, but these local neighbourhoods are likely to be different within different regions of the tissue, we further explored the link with tissue architecture. For dataset 1 we also had information about the three tissue domains that each cell had been assigned to during image segmentation, i.e., tumor, normal and interface (*9*). As the communities had a non-random distribution across the domains, we visualised the distribution of each community relative to the cross section through the tissue, to further expand on this spatial organisation of the communities (Fig. 2a). For example, in the Vehicle setting, community 18, with high endothelium and B cell portion was restricted to the normal non-tumor region as its cell count diminished going into the tumor bulk (Supp. Fig. 2a). Additionally, community 10, with high Type 1 macrophage contribution peaked in cell count at the tumor boundary, demonstrating a clear interface region between normal tissue and the tumor bulk (Supp. Fig. 2b). There were also further communities, number 2 and 3, that were found only in the tumor region, albeit at very different frequencies between Vehicle and MRTX1257 conditions (Supp. Fig. 2c, d). We have previously observed that following treatment with MRTX1257, the immune excluded phenotype of this tumor model was remodelled into a more inflammatory immune infiltrated TME (*9*). Here, we could see that this conversion was also reflected in spatial distribution of the communities, suggesting that tissue domain definition was lost following KRAS-G12C inhibition. Not only did the concentration of communities such as 10 and 18 to the interface and normal regions become less pronounced, but also spatial patterns within the tumor bulk changed following KRAS-G12C inhibition, as the frequency of many communities, such as 2 (type 2 macrophage dominant), 3 (tumour dominant) and 16 (mixed phenotype with high type 2 macrophage portion) were altered, suggesting transition to a new organisation of the TME (Fig. 2b, Supp. Fig. 2e).

**Figure 2.**
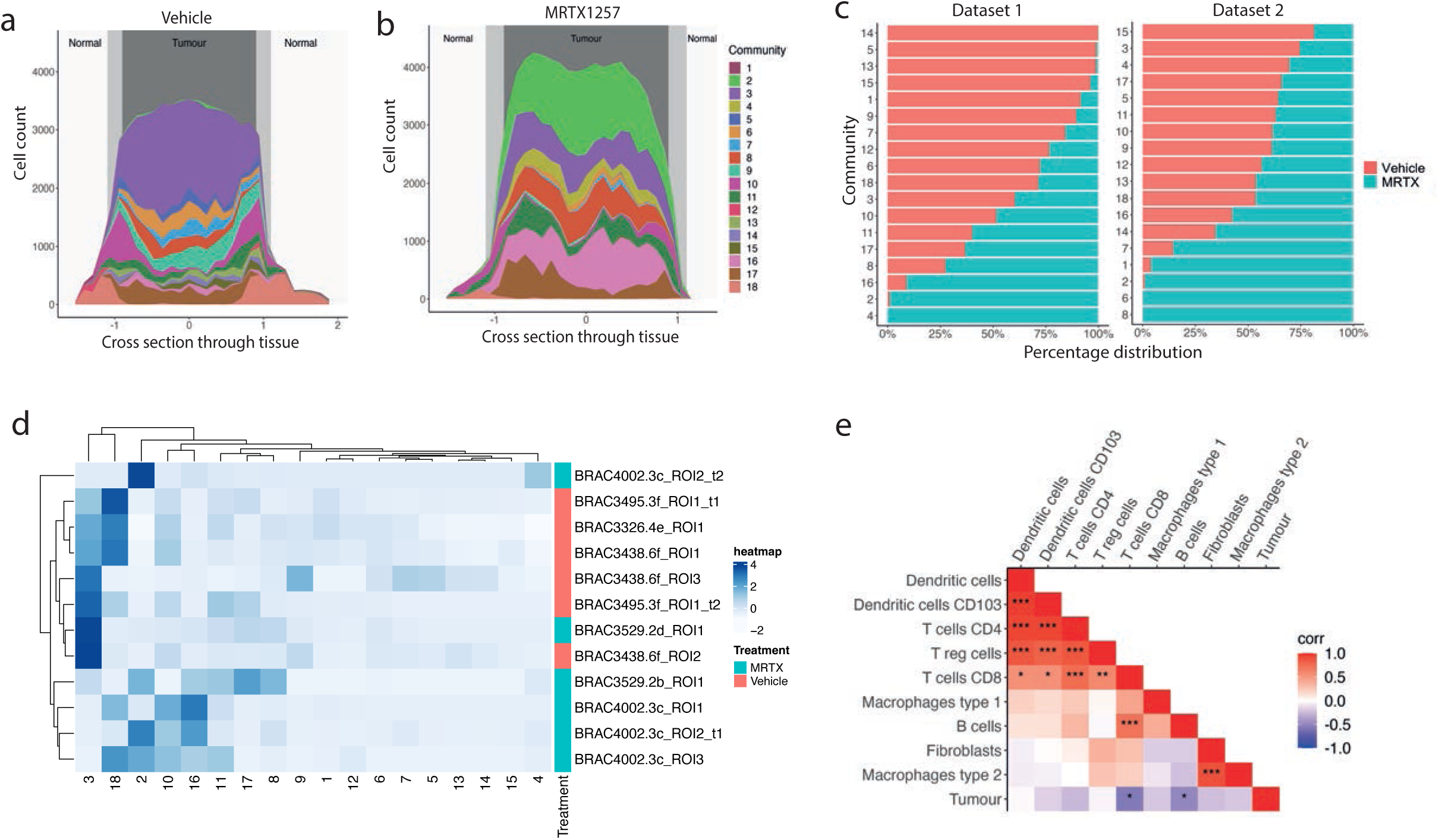
Spatial communities reflect refined tissue architecture and have functional relevance. **a)** Cell count per community relative to cross section through the tissue, where 0 represents the centre point of the tumor in **a)** Vehicle and **b)** MRTX1257 treatment settings. **c)** Percentage distribution of each community across Vehicle (left) and MRTX1257 (right) treatment groups. Bars ordered by increasing percentage distribution in Vehicle setting. **d)** Hierarchical clustering of community proportion per ROI for dataset 1, with use of dendrograms to show relationships between similar ROIs, similar communities, and community distribution across the treatment groups. **e)** Pearson correlation calculation on the proportion of each cell type pair within each community * = p < 0.05, ** = p < 0.01, *** = p < 0.001. Cell types clustered based on correlation value.

Comparing the relative contribution of each community between the Vehicle and MRTX1257 treatment groups revealed that prevalence of many spatial patterns was altered following KRAS-G12C inhibition. Some of these shifts in communities captured changes that we have previously described at the single cell level (*9*). For example, communities 5, 13, 14, 15 from dataset 1 were found solely in tumors treated with Vehicle. These communities represent abundant interactions between tumor cells and neutrophils, which were lost or dispersed following treatment with MRTX1257 (Fig. 2c, Supp. Fig. 1d). This is in line with a previous observation that the number of neutrophils within the tumor domain was significantly reduced following treatment (*9*). Alternatively, communities 2, 4 and 16 from dataset 1 and communities 2, 6 and 8 from dataset 2 were almost exclusively found in tumors following treatment with MRTX1257. A shared feature for these communities was a neighbourhood involving high numbers of Type 2 macrophages (F4/80^+^ CD206^+^), which we have previously shown to become greatly increased in number upon KRAS-G12C inhibition (*9*). This agreement with previous observations suggests that the communities are reflecting relevant biological processes. Further supporting this notion was the ability to largely separate tissues into the two treatment groups when clustering the frequency of each spatial community per region of interest (ROI) (Fig. 2d, Supp. Fig. 2f).

One way to infer potential cellular relationships is by correlating cell frequencies, measuring co-occurrences. Therefore, we calculated the correlation of each cell type pair within communities. Strong positive correlations were seen between certain cell types, such as T cells amongst themselves, T cells with dendritic cells, Type 2 macrophages with fibroblasts and CD8^+^ T cells with B cells when calculated per community (Fig. 2e, Supp. Fig. 2g). By contrast, a number of these, such as Tregs with CD4^+^and CD8^+^ T cells, and most T cell-dendritic cell relationships showed lack of significance when quantified in the ROIs (Supp. Fig. 2h, i). This demonstrates the benefit of studying cellular relationships through the identification of localised spatial patterns, as they provide increased statistical power about the interactions occurring in the TME in comparison to measuring interaction of cells across a whole tissue.

### Spatial communities that are abundant in CD8^+^ T cells

Because the abundance of CD8^+^ T cells is associated with positive outcomes in relation to anti-tumoral immune response, we decided to investigate the T cell-rich communities from this tumor model. Previous analysis revealed increased numbers of CD8^+^ T cells inside the tumor domain following KRAS-G12C inhibition in this model (*9*).

The top five communities with the highest CD8^+^ T cell count were identified from dataset 1 and dataset 2 (Supp. Fig. 3a, b). These five CD8^+^ T cell rich communities from both datasets paired up phenotypically, in particular when only shared cell types between datasets were considered (Supp. Fig. 3c). Therefore, unique colours and the names T cell/normal adjacent community (T/NA), T cell/Type 1 macrophage community (T/M1), T cell/Dendritic cell community (T/DC), T cell/Type 2 macrophage community 1 (T/M2_1) and T cell/Type 2 macrophage community 2 (T/M2_2) were assigned to each pair for parallel analysis going forward (Fig. 3a). These 5 communities contained over 75% of the total CD8^+^ T cell population from both datasets, suggesting a representative population of the overall cohort (Supp. Fig. 3d). Interestingly these communities had widely different compositions, placing the T cells in very diverging spatial contexts (Fig. 3a). The T/NA community was characterised by high endothelial cell content, in the T/M1 community the Type 1 macrophages (CD11c^+^ CD68^+^)(*9*) were the most abundant cell type, the T/DC community was strongly enriched in various T cell subsets as well as dendritic cells, and T/M2_1 and T/M2_2 communities were dominated by tumor cells and Type 2 macrophages (F4/80^+^ CD206^+^)(*9*) in different ratios.

**Figure 3.**
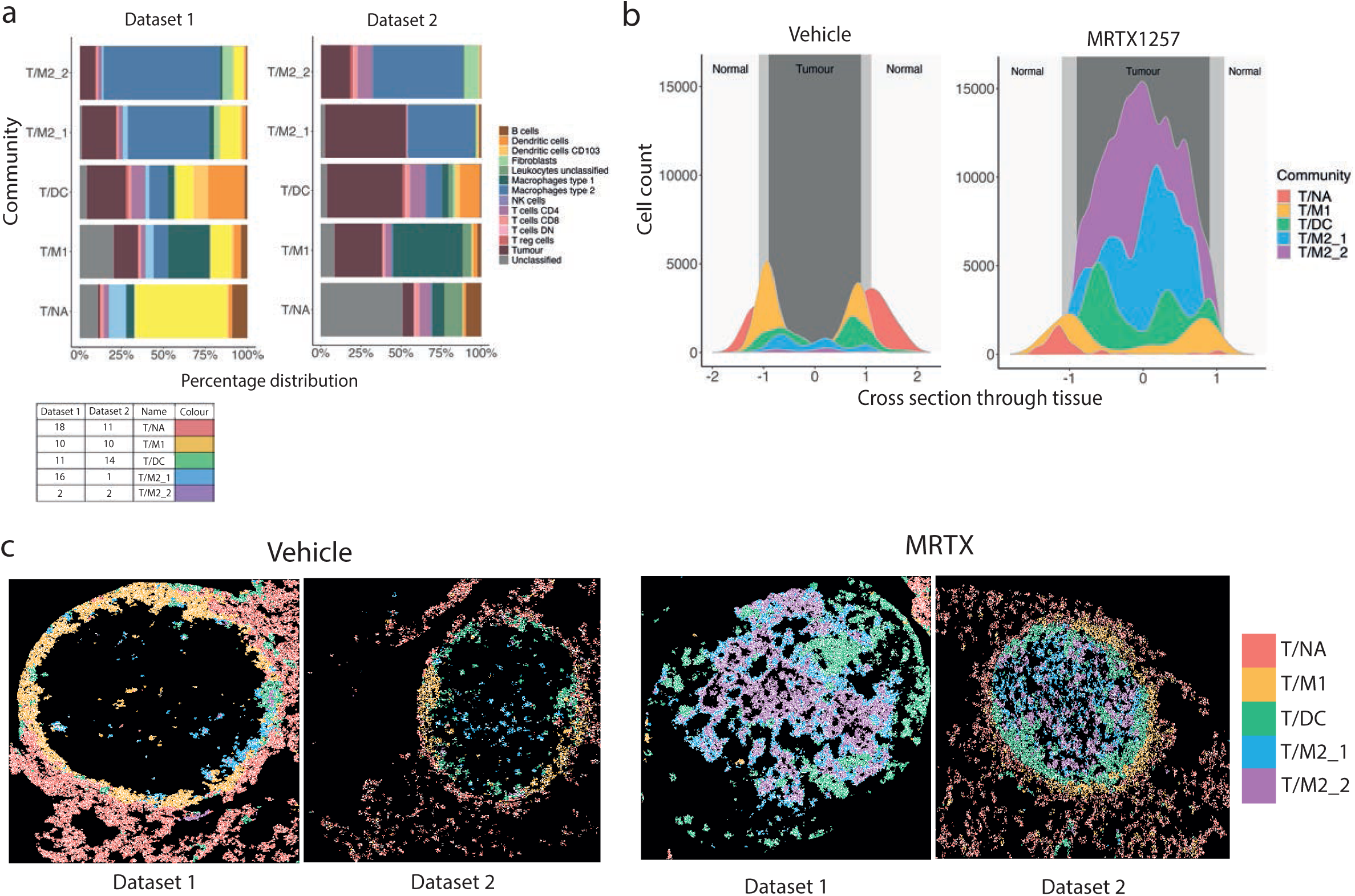
Spatial communities that are abundant in CD8+ T cells. **a)** Percentage distribution of cell types contributing to the top 5 communities with the highest CD8^+^ T cell count from dataset 1 (left) and dataset 2 (right), ordered in pairs based on proportions of shared cell types and labelled with the names T cell/normal adjacent community (T/NA), T cell/Type 1 macrophage community (T/M1), T cell/Dendritic cell community (T/DC), T cell/Type 2 macrophage community 1 (T/M2_1) and T cell/Type 2 macrophage community 2 (T/M2_2). **b)** Cell count of each of the top 5 communities relative to the cross section through the tissue, where 0 represents the centre point of the tumor **c)** Visualisation of cell outlines from cells assigned to T/NA, T/M1, T/DC, T/M2_1 and T/M2_2 communities, with each outline filled with a colour, associating it to one of the five communities, in Vehicle and MRTX1257-treated tumors for datasets 1 and 2.

These CD8^+^ T cell rich communities also differed in presence between treatment groups. The T/NA community was more frequently found following treatment with Vehicle, while the T/M2_2 community was almost exclusively detected in MRTX1257 treated tissues (Supp. Fig. 3e). Following from this, the majority of CD8^+^ T cells in the Vehicle treated tumors were found within T/NA and T/M1 communities, while CD8^+^ T cells in the MRTX1257 treatment group were more likely to reside within T/DC, T/M2_1 and T/M2_2 communities (Supp. Fig. 3f).

Furthermore, the spatial location of the top 5 communities also varied largely in relation to three assigned tissue domains: normal, interface and tumor (Fig. 3b and 3c). In particular, in the vehicle setting, the T/NA community was situated predominantly in the ‘normal’ non-tumor region and the T/M1 community restricted to the ‘interface’, situated as a clear ring around the tumor bulk (Supp. Fig. 3g). The T/DC community was located just inside the tumor bulk in the Vehicle setting but increased in size and its position moved towards the tumor core following treatment with MRTX1257. The T/M2_1 and T/M2_2 communities were concentrated within the tumor domain and predominantly found within the MRTX-treated tumors. Evidently, treatment with MRTX1257 led to a shift in spatial distribution and neighbourhood environment of the CD8^+^ T cells.

### T-cell rich communities respond differently to KRAS-G12C inhibition

Communities were defined based on cell types using lineage markers independently of maturation or activation markers. We next sought to explore how cells within the communities responded to KRAS-G12C inhibition based on the maturation and activation markers included in both datasets. Previously we described how the two subsets of macrophages in this tumor model differed in spatial location and response to treatment with MRTX1257 (*9*). The most notable observation was that the type 2 macrophages increased in cell number and upregulated activation markers such as PD-L1 and MHC-II. Zooming in on the communities here revealed that the upregulated expression could be largely attributed to the type 2 macrophages in the T/DC community (Supp. Fig. 4a and b).

Similarly, we looked at expression of PD-L1 and co-stimulatory receptor CD86 on dendritic cells. Differences were more noticeable across the communities than between treatment groups, suggesting dendritic cell phenotypes were more influenced by surrounding neighbours than treatment with MRTX1257 (Fig. 4a and b, Supp. Fig. 4c). The behaviour of expression varied slightly across these markers, with T/M1 community harbouring high PD-L1 but low CD86 expression in the Vehicle setting, but less so in inhibitor treated conditions. Such high PD-L1 and low CD86 expression is thought to be a characteristic of regulatory or migratory tolerogenic DCs, with a potential to inhibit immune responses (*15*). Alternatively, in T/DC, T/M2_1 and T/M2_2, increased PD-L1 expression on dendritic cells in the MRTX1257 treatment group was accompanied by high CD86 expression, indicative of a more activated DC phenotype making these cells better equipped to facilitate T cell activation. This differential in activation status was further supported by the expression levels of MHC-II, being highest in the T/DC community (Supp. Fig. 4d). Additionally, MRTX1257 treatment increased expression of CXCL9, with the highest overall expression in the T/DC community, suggesting most of the T cell attraction was occurring within this environment (Fig. 4c). This agrees with our finding that the T/DC community contains the largest T cell density. We also compared proliferative marker Ki67 and apoptotic marker cleaved-caspase-3 (c-casp3) expression on tumor cells in the tumor-associated communities (T/DC, T/M2_1 and T/M2_2) and found highest expression of both markers in the T/DC community (Fig. 4d, e, Supp. Fig. 4e, f). This demonstrates the high level of activity occurring within this community, where some tumor cells were thriving whilst others were dying.

**Figure 4.**
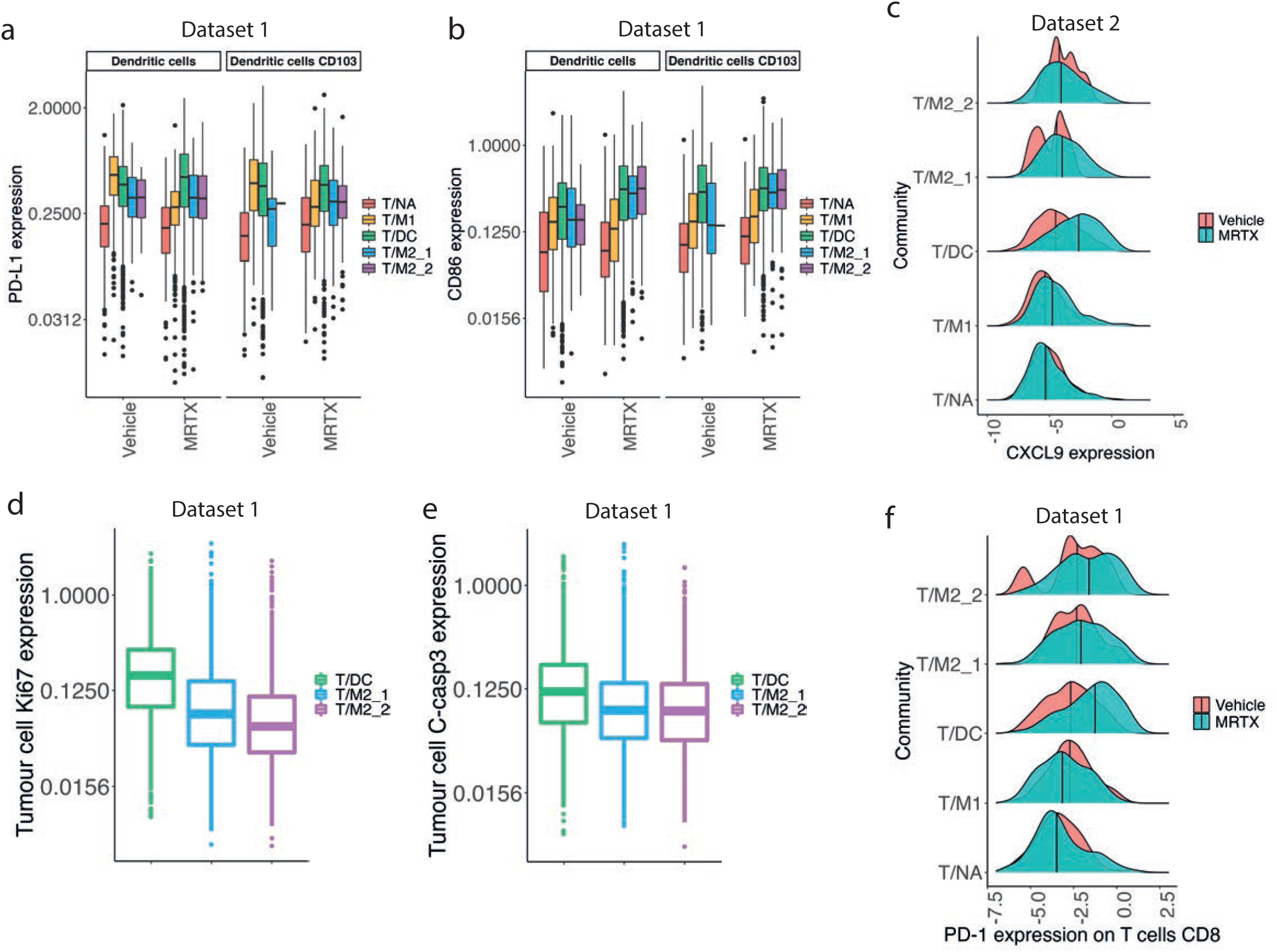
T-cell rich communities respond differently to KRAS-G12C inhibition. **a)** Mean expression of **a)** PD-L1 and **b)** CD86 on dendritic cells and CD103^+^ dendritic cells for T/NA, T/M1, T/DC, T/M2_1 and T/M2_2 communities and Vehicle and MRTX1257 treatment groups for dataset 1 only. Values were log2 scaled. **c)** Mean expression of CXCL9 on dendritic cells, CD103^+^ dendritic cells, macrophages type 1 and macrophages type 2 combined for communities A-E in Vehicle and MRTX1257-treated groups in dataset 2, values were log2 scaled. Centre line shows median expression for each treatment group. **d)** Mean expression of **d)** Ki67 and **e)** cleaved-caspase 3 (c-casp3) on tumor cells in T/DC, T/M2_1 and T/M2_2 communities following treatment with MRTX1257 for dataset 1, values were log2 scaled. **f)** Mean expression of PD-1 on CD8^+^ T cells in communities T/NA, T/M1, T/DC, T/M2_1 and T/M2_2 in Vehicle and MRTX1257-treated groups for dataset 1, values were log2 scaled. Centre line shows median expression for each treatment group.

We expected that these different neighbourhoods would also have an impact on the phenotype of the T cells, so investigated their PD-1 expression to understand how T cell activation state changed following KRAS-G12C inhibition, per community. In the T/NA and T/M1 communities, the PD-1 expression was negligible in both treatment groups, compared to the T/DC, T/M2_1 and T/M2_2 communities, where an increase occurred following treatment with MRTX1257, most pronounced in the T/DC community, indicating a switch in cell state from naïve to activated in these communities following KRAS-G12C inhibition (Fig. 4f, Supp. Fig. 4g). A similar pattern could be seen for CD4^+^ T cells, whereas the T reg cells or ‘Tregs’ had higher PD-1 expression mainly in the T/DC community following MRTX1257 treatment (Supp. Fig. 4h, i). However, expression of LAG-3 was also distinctly higher in CD8^+^ T cells, specifically in the tumor-associated communities, in which ∼30% of PD-1^+^ T cells were also LAG3^+^ following treatment with MRTX. This suggests that activation of the T cells was in part accompanied with induction of T cell exhaustion following KRAS-G12C inhibition (Supp. Fig. 4j, k).

### Positive and negative regulation of anti-tumoral immune responses meet in the T/DC community

While the T/DC, T/M2_1 and T/M2_2 communities contained most of the CD8^+^ T cells in the tumor tissue, these cells expressed significant levels of potential exhaustion markers such as PD-1 and LAG-3. Co-localisation of PD-L1 expressing macrophages and PD-1^+^ CD8^+^ T cells has recently been highlighted as associated with good response to ICI (*16, 17*).

However, previous attempts to reinvigorate these T cells using MRTX1257 in combination with ICIs anti-PD-1 or anti-PD-L1 and anti-LAG3 had failed to achieve any improved tumor control in our 3LL model (*8*). We therefore wondered whether we could gain any insight into the signals provided to the T cells prior to moving into the core of the tumor and becoming incapacitated by exhaustion. The high proportion of activated dendritic cells and increased expression of markers associated with T cell attraction and activation following KRAS-G12C inhibition, as well as evidence for local tumor cell death, pointed towards the T/DC community as a potential cytotoxic T cell activation hub.

As noted, CXCL9 expression was highest within antigen presenting cells in the T/DC community (Fig. 4c). Separating dendritic cells based on their CXCL9 expression into ‘CXCL9 low’ and ‘CXCL9 high’ groups showed that CXCL9 high dendritic cells had a significantly shorter distance to their nearest PD-1^+^ CD8^+^ T cell, PD-1^+^ CD4^+^ T cells and PD-1^+^ Tregs (Fig. 5a and Supp. Fig. 5a, b). Visual inspection indeed confirmed that CXCL9 high dendritic cells were frequently found in proximity to and interacting with PD-1^+^ CD8^+^ T cells (Fig. 5b). CXCL9 expression could therefore be one of the key mediators driving the aggregation of the T cells and dendritic cells within the T/DC community, as part of a mechanism to draw in the activated T cells to launch an anti-tumor immune response. This would be in agreement with our previous work showing that CXCL9 expression was one of the strongest predictors of ICI (*18*).

**Figure 5.**
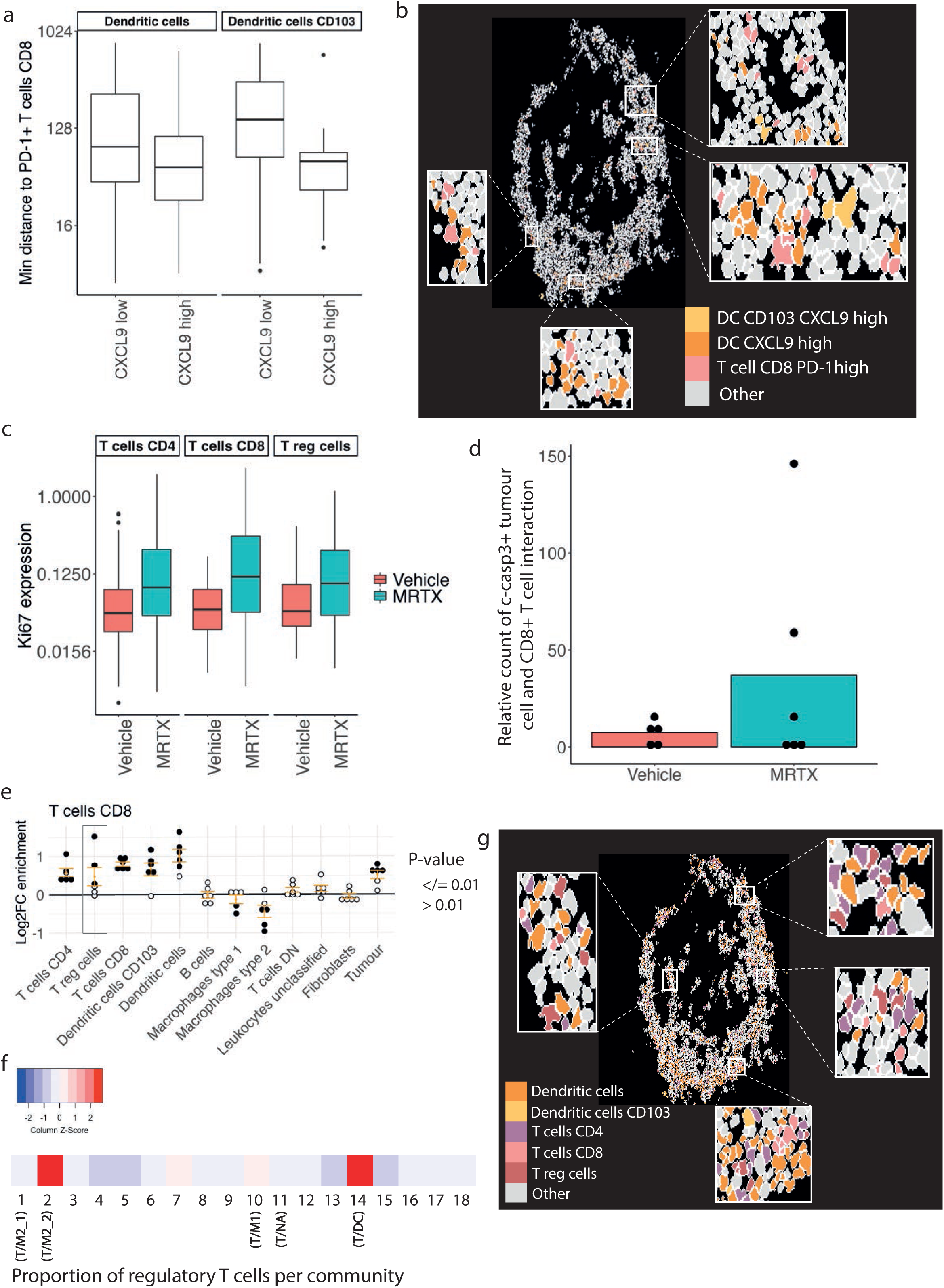
Positive and negative regulation of anti-tumoral immune responses come together in the T/DC community. **a)** Minimum distance of dendritic cells and CD103^+^ dendritic cells that have ‘low’ or ‘high’ CXCL9 expression (based on a threshold of 0.5) to PD-1^+^ CD8^+^ T cells within 800 pixels in the T/DC community from dataset 2, distance values were log2 scaled. **b)** Visualisation of cell outlines for cells assigned to the T/DC community in MRTX1257-treated tissues from dataset 2, with CXCL9 high dendritic cells and CD103^+^ dendritic cells, and PD^+^1 CD8^+^ T cells coloured in to show spatial proximity of these cell phenotypes. Some regions have been expanded for easier visualisation. **c)** Mean expression of Ki67 on CD4^+^, CD8^+^ and regulatory T cells within the T/DC community for Vehicle and MRTX1257 treatment groups from dataset 2, values were log2 scaled. **d)** Number of times a c-casp3^+^ tumor cell is found in the 15-pixel neighbourhood of a CD8^+^ T cell within the T/DC community, compared across Vehicle and MRTX1257 treatment groups for dataset 2, averaged per ROI. Count is relative to the proportion of tumor cells that were c-casp3^+^ in Vehicle vs MRTX1257 treatment groups. Each dot represents the value of one ROI. **e)** Log2 fold changes in enrichment from neighbouRhood analysis for CD8^+^ T cells in the T/DC community following treatment with MRTX1257. Filled circles represent images from which enrichment value was statistically significant compared to randomisation of the spatial arrangements following treatment with MRTX1257 for dataset 2. **f)** Scaled proportion of regulatory T cells contributing to each of the 18 original communities for dataset 2. **g)** Visualisation of cell outlines for cells assigned to the T/DC community in MRTX1257-treated tissues for dataset 2, with dendritic cells, CD103^+^ dendritic cells and CD4^+^, CD8^+^ and regulatory T cells filled in to illustrate spatial proximity of these cell types. Some regions have been expanded for easier visualisation.

We sought to determine further indication of active anti-tumoral immune response within this community. A significant increase of Ki67 expression was identified for CD4^+^ and CD8^+^ T cells on MRTX1257 treatment, and a similar trend for Tregs, suggesting that the T cells had increased proliferation following KRAS-G12C inhibition (Fig. 5c). Frequency of casp3^+^ tumor cells was highest in the T/DC community and the occurrences in which a c-casp3^+^ tumor cell was found in the 15-pixel neighbourhood of a CD8^+^ T cell increased following treatment with MRTX1257 (Fig. 5d). These interactions were primarily found within the T/DC community (Supp. Fig. 5c). There was no such increase in spatial interactions identified for CD4^+^ or Tregs with c-casp3^+^ tumor cells (Supp. Fig. 5d). Such increased interactions point to increased cytotoxicity of the T cells against the tumor cells following KRAS-G12C inhibition, suggesting that the neighbourhood of the T/DC community was indeed likely to be able to support the effector function of the CD8^+^ T cells.

We then wondered why such a cytotoxic response was not effective enough to mediate clinical benefit and what could be driving the induction of T cell exhaustion or dysfunction. To identify potential negative regulatory influences of the immune response, we employed another way of interrogating spatial relationship, by calculating enrichment scores for cells in the near neighbourhood, compared to randomised data (*9, 19*). Following MRTX1257 treatment, it was determined that CD4^+^ T cells and dendritic cells were significantly enriched in the neighbourhood of a CD8^+^ T cell within the T/DC community in at least five out of the six images, compared to random permutation (Fig. 5e). This is supportive of a microenvironment promoting anti-tumoral immune response, which was not apparent from T/NA, T/M1, T/M2_1 and T/M2_2 communities (Supp. Fig. 5e-h). However, Tregs were also significantly enriched in the neighbourhood of CD8^+^ T cells in 3 out of the 6 images, indicating the presence of immune suppressor cells in the vicinity where T cell activation and effector functions may be taking place (Fig. 5e). Treg frequencies were highest within the T/DC and T/M2_2 communities (Fig. 5f, Supp. Fig. 5i). This T cell subset also increased in size, most notably in the T/DC and T/M2_2 communities, and as we saw previously, was showing evidence of increased activation after MRTX1257 treatment (Supp. Fig. 5j, Supp. Fig. 4i). Indeed, Tregs were seen intermixed with effector T cells and dendritic cells, suggesting a potential role in locally dampening anti-tumoral immune responses (Fig. 5g).

### Assigning an important role for Tregs in dampening anti-tumoral immune responses

Upon revealing the high presence of Treg cells within the T/DC community and showing that they are enriched within the neighbourhood of CD8^+^ T cells and neighbouring dendritic cells and CD4^+^ T cells following treatment with MRTX1257, we decided to explore their role within this community in relation to anti-tumoral immune response. We therefore split up the T/DC community into two neighbourhoods: those with presence of Treg cells, named ‘Tregs’, or those with absence of Treg cells, named ‘No Tregs’. The neighbourhoods differed slightly in their composition of cell types, with the ‘Tregs’ neighbourhood comprising a higher proportion of dendritic cells and CD4^+^ T cells, whereas the ‘No Tregs’ neighbourhood contained a higher tumor and type 2 macrophage portion (Fig. 6a).

**Figure 6.**
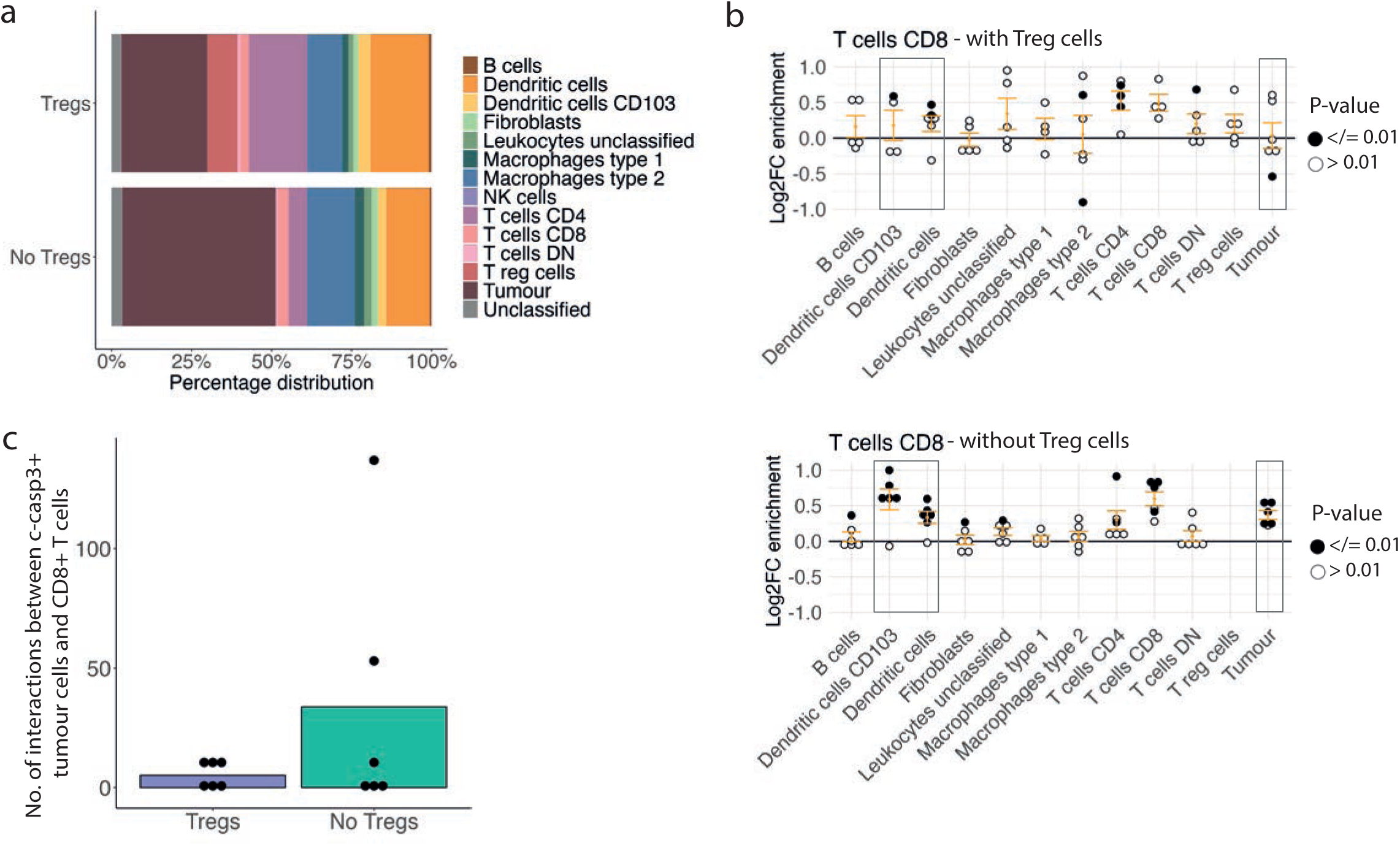
Assigning an important role for Tregs in dampening anti-tumoral immune responses. **a)** Percentage distribution of cell types contributing to ‘Tregs’ and ‘No Tregs’ neighbourhoods within the T/DC community following treatment with MRTX1257. **b)** Log2 fold changes in enrichment from neighbouRhood analysis for CD8^+^ T cells in ‘Tregs’ (top) and ‘No Tregs’ (bottom) neighbourhoods within the T/DC community following treatment with MRTX1257. Filled circles represent images from which enrichment value was statistically significant compared to randomisation of the spatial arrangements within the T/DC community following treatment with MRTX1257 for dataset 2. **c)** Number of times a c-casp3^+^ tumor cell is found in the 15-pixel neighbourhood of a CD8^+^ T cell within the T/DC community, compared across ‘Tregs’ and ‘No Tregs’ neighbourhoods in dataset 2, averaged per ROI. Count is relative to the proportion of tumor cells that were c-casp3^+^ in ‘Treg’ vs ‘No Treg’ groups.

While the frequency of CD8^+^ T cells within both the ‘Tregs’ and ‘No Tregs’ neighbourhoods were similar (Supp. Fig. 6a), the cellular interactions for CD8^+^ T cells differed substantially between these environments. Despite the slightly higher dendritic cell frequency in the ‘Tregs’ neighbourhoods, we saw a lack of spatial enrichment between CD8^+^ T cells and both dendritic cell subsets when Treg cells were present, in contrast to a positive spatial enrichment of dendritic cells in the CD8+ T cell neighbourhood when Treg cells were absent (Fig. 6b, Supp. Fig. 6b). Additionally, when Tregs were not present in the CD8^+^ T cell neighbourhood, a strong enrichment of tumor cells was identified following MRTX1257 treatment, compared to slight depletion when the Tregs were nearby. Furthermore, the frequency of c-casp3^+^ tumor cells in the CD8^+^ T cell close neighbourhood increased on MRTX1257 treatment when Treg cells were absent (Fig. 6c). We therefore only saw interactions with CD8^+^ T cells indicative of an active anti-tumoral immune response when the Treg cells were absent, suggesting an inhibitory role for the Treg cells. These changes in neighbourhood enrichment were not observed when comparing CD4^+^ T cells with or without Treg cells in their neighbourhood, indicating that Treg presence may affect CD8^+^, but not CD4^+^ T cell relationships (Supp. Figs. 6c and 6d). However, there were several other changes to the cellular interactions when sub-setting the T/DC community based on presence or absence of Tregs, suggesting that Treg cells may be impacting the local milieu of this community in many ways, further pointing towards their potential negative influence on anti-tumoral immune response (Supp. Fig. 6b).

Overall, these analyses indicated that the T/DC community potentially could provide an activating environment for the anti-tumor cytotoxic T cell response, but the presence of Treg cells was likely imposing a strong negative influence.

A similar analysis for the T/M2_2 community, also relatively rich in Tregs (Fig. 5f), showed that CD8^+^ T cell and Treg interactions focussed mainly around fibroblasts, while CD8^+^ T cell interactions with M2 macrophages were largely interrupted in presence of Tregs (Supp. Fig. 6e). Likewise, CD4^+^ T cell interactions with M2 macrophages were also diminished in presence of Tregs (Supp. Fig. 6f). This further solidifies the likely role of Tregs suppressing T cell function through downregulating interactions with antigen presenting cells following KRAS-G12C inhibition, which occurred across multiple spatial groups.

### Treg spatial communities in human lung non-small cell lung cancer

While community analysis in mice can help us to identify recurring patterns in a fairly homogeneous and controlled experimental setting, we ultimately aim to translate our findings to better treat patients. Therefore, we wanted to investigate whether communities similar to the mouse communities could be identified in human lung cancer clinical samples, focussing on the communities rich in Tregs, effector T cells and dendritic cells, like our mouse T/DC community.

Recently, 151 tumor regions from 81 untreated NSCLC patients from the TRACERx longitudinal study were analysed with imaging mass cytometry using two 35-plex antibody panels: panel 2, the pan-immune cell panel, and panel 1, designed to assess the differentiation states of T cells and stromal cells in greater detail (T cell panel). The T cell panel provides us with an opportunity to not only match cell type compositions with our mouse data, but also to potentially refine the T cell signatures that are associated with them (*20*). Therefore, we chose to detect communities based on the T cell panel (panel 1) with relatively high granularity by optimising parameters of the Schurch *et al.* method, obtaining 30 communities, including 7 that were rich in Tregs and other T cells (Fig. 7a). Five of these communities (*p1_C7*, *p1_C17*, *p1_C23*, *p1_C26* and *p1_C27*) were including mature and/or exhausted CD4^+^ and CD8^+^ T cells, similar to the mouse T/DC community. Another characteristic feature of the mouse T/DC community was the presence of dendritic cells, as well as the peritumoral localisation. Dendritic cells could not be identified from the T cell panel (panel 1, *p1*). However, community analysis of the pan-immune (panel 2) data had previously identified 10 communities in which *p2_C1: tumor border* also contained dendritic cells and was in the peritumoral region (Supp. Fig. 7a, b, c). Interestingly, the presence of *p2_C1: tumor border* correlated with the density of Tregs in LUAD (Fig. 7b), but not in LUSC (Supp. Fig. 7d). In a major subset of patients, we could confirm the co-occurrence of *p2_C1: tumor border* from panel 2 and Treg communities from panel 1 (Fig. 7c), and that in particular Treg communities *p1_C7*, *p1_C17*, *p1_C23*, *p1_C26* and *p1_C27* displayed a peritumoral distribution close to *p2_C1: tumor border* (Fig. 7d, e). Homing in on the phenotypes of the T cells associated with these communities revealed that *p1_C27* was standing out among the others. CD4^+^and CD8^+^ T cells in *p1_C27* expressed high levels of immune checkpoints (TIM3, GITR, CTLA-4, ICOS). However, in contrast to the mouse T/DC community, there was no evidence of ongoing T cell activation, with absence of proliferation marker Ki67 and cytotoxicity marker Granzyme B (GZMB) (Fig. 7f). The Tregs in *p1_C27* were highly expressed all tested markers except GITR, an immune checkpoint that was recently described to mark the most immune suppressive Treg subset in NSCLC and associated with PD-1 resistance (*21*). Some of the other communities expressed similar or higher levels of GITR on the Tregs (*p1_C26*, *p1_C23*, *p1_C6*, *p1_C17*) and showed above average expression of immune checkpoints on the CD4^+^and CD8^+^ T cells (*p1_C7*, *p1_C23*, *p1_C17*, *p1_C26)*, more similar to the mouse T/DC community. The MYSTIC trial (*22*) demonstrated that the combination of durvalumab (anti-PD-L1) and tremelimumab (anti-CTLA-4) only showed improved survival compared to durvalumab alone in patients with high TMB (>20 mutations/Mb). Using the genomics data available for the patients of this TRACERx cohort (*23*), we found that *p1_C7* and *p1_C17* also correlated with the most increased TMB in LUAD, suggesting that targeting of these Treg communities with anti-CTLA-4 might be beneficial (Fig. 7g). Overall, these data from human lung cancer samples that have not been treated with KRAS inhibitory drugs suggest that cellular communities similar to the mouse T/DC community exist at baseline, where CD4^+^and CD8^+^ T cells, dendritic cells and Tregs were gathered together at the tumor periphery.

**Figure 7.**
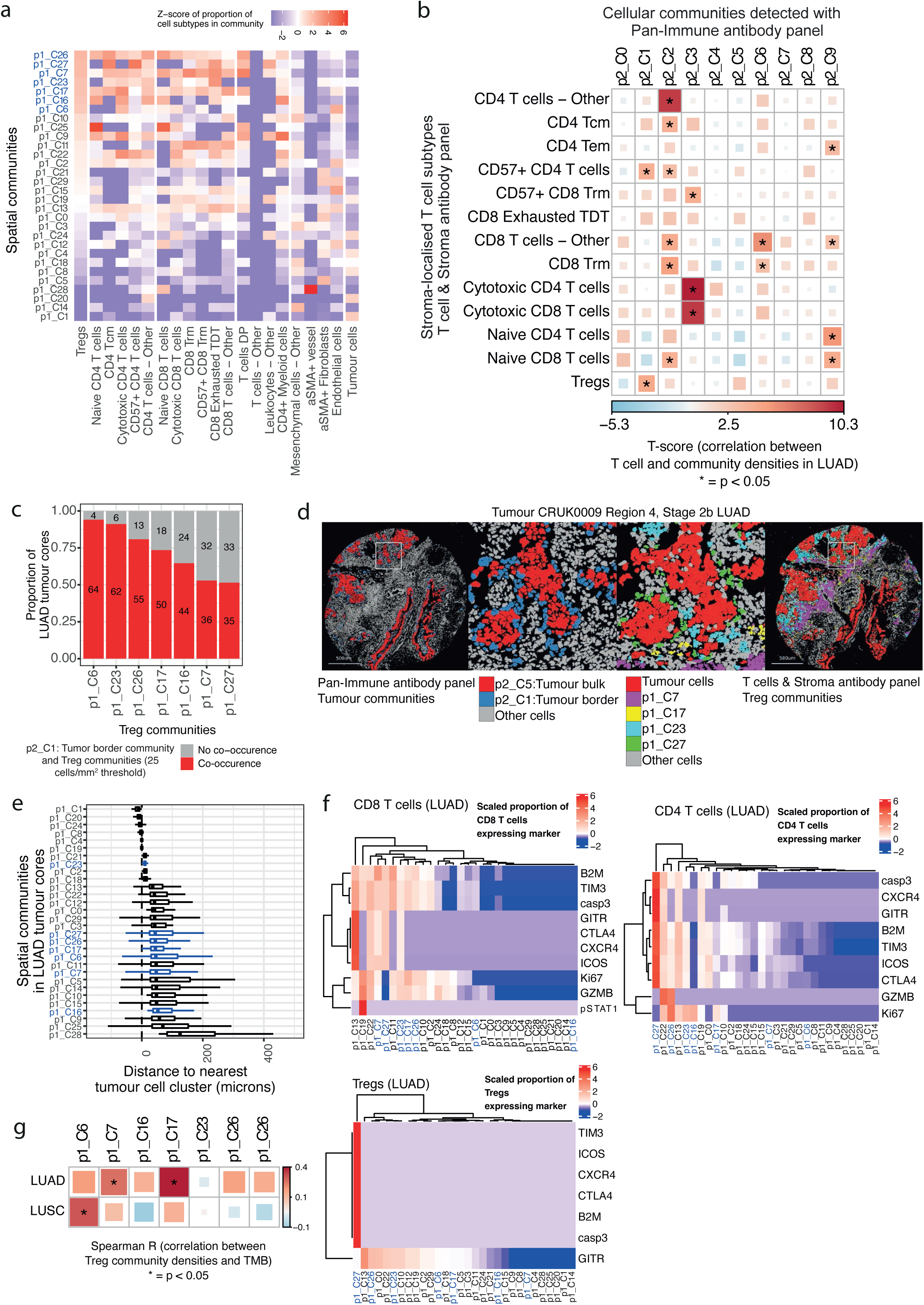
Treg spatial communities in human lung non-small cell lung cancer. **a)** Thirty spatial communities detected in 137 tumor cores from 70 patients with non-small cell lung cancer. The Z-score of the proportion of cell subtypes, detected using the T cell & Stroma antibody panel (*20*) in each spatial community is shown. Communities *p1_C6*, *p1_C7*, *p1_C16*, *p1_C17*, *p1_C23*, *p1_C26*, *p1_C27* (blue lettering) contain the highest proportions of Tregs. **b)** Tregs and CD57^+^ CD4^+^ T cells are associated with *p2_C1: Tumor border* spatial communities. Correlation between the density of stroma-localised T cell subtypes detected using the T cells & Stroma antibody panel and the cell density of ten spatial communities detected using the Pan-immune antibody panel, in 68 LUAD tumor cores from 40 patients. Linear mixed effects model (LMEM) to test the association between T cell density and cell density of communities with patient as a random covariate to adjust for multiple cores per tumor. Analysis of variance test comparing the LMEM to the null model, p-values are unadjusted. *:p < 0.05. **c)** Proportion of LUAD tumor cores that contain at least 25 cells/mm^2^ of Treg communities (*p1_C6*, *p1_C7*, *p1_C16*, *p1_C17*, *p1_C23*, *p1_C27*) and *p2_C1: Tumor border* communities. 68 LUAD tumor cores from 40 patients. **d)** Pseudo-coloured images highlighting cells in the *p2_C1: Tumor border* communities corresponding to cells in Treg communities in serial tumor cores. **e)** Per-image median distance between cells of a community and their nearest tumor cell cluster. Tumor cell clustering method described in Magness *et al.,* manuscript in review. 72 LUAD tumor cores from 48 patients. **f)** Heatmap displaying the scaled proportion of CD8^+^ T cells, CD4^+^ T cells and Treg cells expressing phenotypes of interest. 72 LUAD tumor cores from 48 patients. **g)** Spearman correlation between the density of Treg communities and total harmonised tumor mutational burden. 70 LUAD cores from 41 patients, 49 LUSC cores from 21 patients.

### Assigning an important role for Tregs in dampening anti-tumoral immune responses

We then went back to the mouse model to investigate whether depleting Treg cells using an Fc-optimised anti-CTLA-4 antibody (*24*) could promote immune mediated control of orthotopic 3LL ΔNRAS lung tumors treated with KRAS-G12C inhibitor. Anti-CTLA-4 antibody along with anti-PD-1 therapy failed to control tumor growth and extend survival of mice. Similar to what we observed previously, MRTX1257+anti-PD-1 led to short-term tumor control after one week on treatment and extended survival, but eventually tumors relapsed and nearly all mice reached their endpoint within 5-6 weeks (*8*) (Fig. 8a, b). Our previous observations showed that anti-PD-1 antibodies did not improve the response to MRTX1257 in this system (*8*). Also, after one week of treatment, combination of MRTX1257+anti-PD-1 with Treg depleting anti-CTLA-4 therapy, did not significantly impact tumor growth (Fig. 8b). However, after two weeks, tumor volume assessment revealed most tumors from MRTX1257+anti-PD-1 group had relapsed in contrast to triple combination with anti-CTLA-4 where tumors kept regressing (Fig. 8c, Supp. Fig. 8a). This response pattern continued after 3 weeks of treatment (Supp. Fig. 8b), and the effect of the triple combination in the long-term resulted in significant tumor control and extension of survival compared to MRTX1257+anti-PD-1, even generating 1/8 completely tumor free mice, which remained tumour free after withdrawal of treatment for 30 days (Fig. 8a).

**Figure 8.**
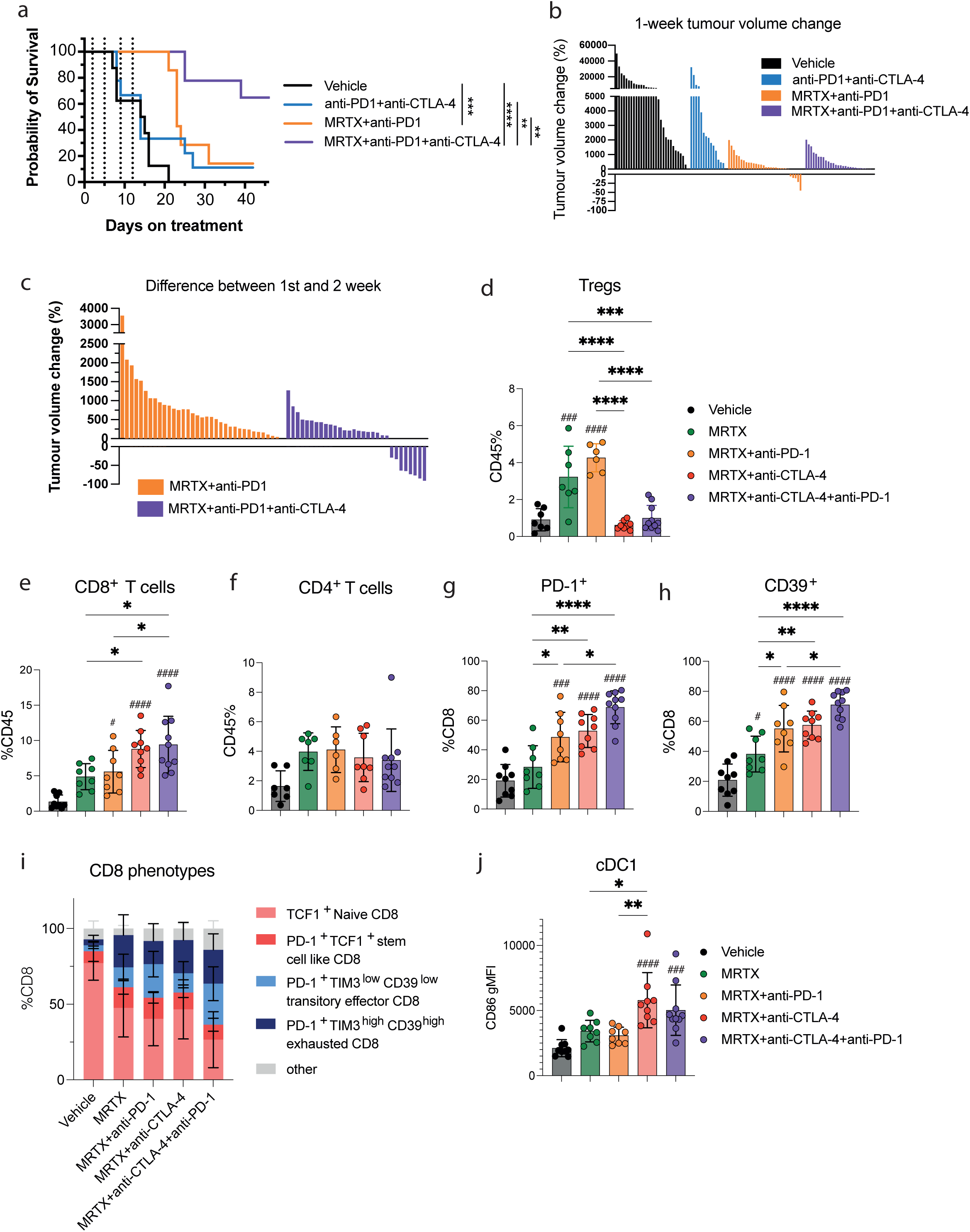
Assigning an important role for Tregs in dampening anti-tumoral immune responses. **a)** Kaplan-Meier analysis of the overall survival of mice using the 3 Lewis Lung carcinoma model under Vehicle, anti-PD-1+anti-CTLA-4, MRTX1257+ anti-PD-1 and MRTX1257+anti-PD-1+anti-anti-CTLA-4 treatment groups. b) Tumor volume changes after 1 week of treatment as measured by μCT scanning. c) Tumor volume changes after the second week of treatment as measured by μCT scanning, for MRTX1257+anti-PD-1 and MRTX1257+anti-PD-1+anti-CTLA-4 treatment groups. d) Percentage of all CD45^+^ cells identified as T regulatory cells (gated as CD45^+^ CD3^+^ CD4^+^ Foxp3^+^) measured by flow cytometry in the tumor (each dot represents a mouse, one-way ANOVA). e) Percentage of all CD45^+^ cells identified as **e)** CD8^+^ T cells (gated as CD45^+^ CD3^+^ CD8^+^) and **f)** CD4^+^ T cells measured by flow cytometry (each dot represents a mouse, one-way ANOVA). g) Percentage of CD8^+^ T cells that are **g)** PD-1^+^, **h)** CD39^+^ measured by flow cytometry in the tumor. i) Distribution of CD8^+^ T cells in the tumor that are TCF1^+^, PD-1^+^TCF1-, PD-1^+^TIM3^low^CD39^low^ and PD-1^+^TIM3^high^CD39^high^ across treatment groups. j) Percentage of DC1s in the tumor that are CD86^+^ measured by flow cytometry.

Flow cytometry analysis demonstrated increased Treg (CD4^+^FOXP3^+^) infiltration upon MRTX1257 treatment, which was exacerbated with the addition of anti-PD-1, while the addition of anti-CTLA-4 therapy induced depletion of Tregs in the tumors, as expected (Fig. 8d). This occurred with an increased CD8^+^ T cell infiltration, beyond what was observed with MRTX1257 alone or in the MRTX1257+anti-PD-1 combination (Fig. 8e), while total CD4^+^ T cell numbers remained stable between all conditions (Fig. 8f). CD8^+^ T cells in the tumor upregulated activation markers PD-1 and CD39, suggestive of increased tumor-reactive T cell rates following Treg depletion (Fig. 8g, h), and progressively converted from predominantly naïve in the vehicle control setting to effector and exhausted CD8^+^ T cells with addition of the three therapeutics (MRTX1257, anti-PD-1 and anti-CTLA-4, Fig. 8i). Moreover, upon combination treatment with MRTX1257+anti-PD-1+anti-CTLA-4, we observed elevated CD86 expression on CD103^+^ Type 1 Dendritic cells (cDC1s), an indication of increased antigen presentation and explainable by reduced transendocytosis in absence of Tregs (Fig. 8j).

Interestingly, in the pulmonary lymph nodes, Tregs were not depleted by the addition of anti-CTLA-4 (Supp. Fig. 8d). This has been observed before and was related to the differential expression of CTLA-4 on Tregs, being high in tumors and low in the lymph nodes (*25, 26*), which is in agreement with our data (Supp. Fig. 8e). Despite this, the lymph nodes showed pronounced T cell activation, evidenced by increased expression of PD-1 and proliferation marker Ki67 in the triple combination treatment group compared to double combinations or single agent MRTX1257 (Supp. Fig. 8f,g), indicative of the priming of a systemic immune response.

## DISCUSSION

Despite objective response rates of roughly 30-40%, the licensed KRAS-G12C inhibitors sotorasib(*3, 27, 28*) and more recently adagrasib(*29, 30*) have so far achieved only a modest improvement in progression free survival of about one month and no improvement in overall survival. This very limited clinical benefit has been attributed to intrinsic and acquired resistance mechanisms(*31–34*), which has led to increased efforts to explore combination therapies. Besides combinatorial targeting of multiple tumor cell intrinsic pathways(*35–38*), early pre-clinical experiments also suggested a potential synergy with immune therapy(*6, 39*). Inhibiting KRAS-G12C was able to turn a cold tumor into a hot pro-inflammatory environment, potentially supporting anti-tumor immune responses in combination with ICI(*9*). However, our own pre-clinical work demonstrated that despite the presence of TME changes, anti-PD-1 refractory tumors did not become responsive to ICI combinations as a result of KRAS-G12C inhibition(*8*). Similarly, first reports of the safety and efficacy of sotorasib in combination with pembrolizumab or atezolizumab in advanced KRAS-G12C NSCLC have suggested poor response rates in patients that previously progressed on ICI, at least in part due to severe combination toxicities (*11*).

To better understand this persisting resistance to therapy, we further investigated our most immune refractory KRAS-G12C mutant NSCLC model. The 3LL Lewis Lung Carcinoma model has a history of repeated passaging through immunocompetent mice and as a result has developed a complex mixture of immune resistance mechanisms (*40*). Indeed, in previous work treating orthotopic 3LL lung tumors with KRAS-G12C inhibitors only provided temporary tumor control, and combinations with anti-PD-1 or anti-PD-L1 and anti-LAG3 failed to give additional benefit(*8*).

Recent studies have demonstrated that the organisation and distribution of cellular communities in the tumor can be predictive of clinical outcome (*14, 41, 42*), including studies on lung cancer (*13, 43*). Similarly, in this study, we investigated the cellular communities in our therapy-resistant 3LL model for evidence of immune suppressive patterns. Similar to the “suppressed expansion” cluster in breast cancer described by Danenberg et al. (*41*), we identified a T cell rich cellular community (T/DC) with evidence of T cell activation, proliferation, exhaustion and inhibition. In this cluster we saw abundant interactions between CD4^+^ and CD8^+^ T cells and mature antigen presenting DCs, similar to interactions normally seen within the T cell zones of lymph nodes, however not resembling tertiary lymphoid structures as B cell involvement was sparse. Furthermore, we found evidence of anti-tumor cytotoxicity within this community, with enrichment for interactions between CD8^+^ PD-1^+^ T cells and cleaved-caspase 3 expressing tumor cells, suggestive of T cell-induced apoptosis. After treatment with KRAS G12C inhibitory drug, we observed increased CXCL9 expression, which is involved in T cell recruitment, potentially from the periphery. Tumor draining lymph nodes (tdLNs) are considered to play a key role in facilitating the anti-tumor immune response, with the priming of new waves of antitumor T cells by both migratory and resident DC in tumor and tdLN, which are subsequently attracted to the tumor by chemokine expressing DC in the TME (*44*). Indeed, recent work showed that patients responding to ICI harboured more shared TCR clones between tumor and tdLNs, but also provided evidence for local expansion and proliferation of CD8^+^ T cells in the TME (*45*). It remains to be determined to what extent the make-up of spatial communities in the TME are a direct reflection of processes occurring in the tdLNs.

While Treg cells were generally quite scarce in the data, they were found strongly enriched within this T cell/DC-enriched community. Tregs have direct and indirect mechanisms to inhibit T cell function. Firstly, Tregs can inhibit CD8^+^ T cells directly by secreting or exposing T cell suppressive cytokines, such as IL-10, IL-35 and TGF-beta, by scavenging IL-2, and by the generation of extracellular adenosine (*46*). Unfortunately, we could not investigate these mechanisms within the spatial communities as our panel lacked the markers for these soluble factors.

Secondly, Tregs can compete with CD8^+^and CD4^+^ T cells for the co-stimulatory molecules on the antigen presenting cells, primarily by transendocytosis or trogocytosis of CD80/CD86 through the interaction with CTLA-4(*47, 48*). Therefore, blocking of CTLA-4 is able to enhance CD8^+^ T cell responses. CD8/DC interactions were indeed less frequently observed in the neighbourhoods containing Tregs, potentially reducing the opportunity for local activation of CD8^+^ T cells. Furthermore, depletion of Tregs using anti-CTLA-4, in combination with MRTX1257 and anti-PD-1, led to increased expression of activation markers, and most importantly, the CTLA-4 ligand CD86 on the DCs and enabled tumor control by enhanced immunity in this otherwise strongly immune evasive model.

This raises the question of whether the presence of a cellular community that is rich in activated T cells and DCs but also Tregs may predict resistance to the combined therapy of KRAS G12C inhibition and PD-1 blockade. Spatial investigation of cellular communities in the pre-treatment biopsies of the CodeBreak 100/101 clinical trials, would allow for correlation with clinical outcome. This is highly relevant as initial results in this study showed that the combination of Sotorasib with PD-1 or PD-L1 led to unexpected liver toxicity(*49*), hence identifying and excluding patients that are resistant to this combination due to Treg activity would help to prevent unnecessary ineffective treatments. Furthermore, this could open up the possibility of using anti-CTLA-4 or other Treg targeting therapies in combination with sotorasib in selected patients based on a spatial Treg community biomarker.

But, as our data suggests, Treg rich communities are not unique to the KRAS inhibitor treated conditions, as the T/DC community was present at similar frequencies in the vehicle setting. Likewise, a deeper analysis of the RUBICON cohort (*20*) demonstrated that various Treg rich communities are present in a significant proportion of treatment-naïve patients. Furthermore, Pentimalli *et al.* described the presence of a niche rich in cytotoxic T cells, regulatory DCs and Tregs, marked by localised expression of CXCL9, similar to our observations, as well as CCL19 and CCL21, in 3D spatial transcriptomics analysis of a high grade NSCLC patient (*50*). This suggests that DC, T cell and Treg rich communities are a recurring feature in lung cancer and may predict response in a broader context. Several studies in lung cancer so far have indicated frequencies of intra-tumoral or circulating Tregs to be associated with poor outcome (*51–53*) or therapy resistance(*54, 55*), but few have looked into the cellular communities in more detail.

Sorin *et al.* identified B cell communities that associated with survival advantage unless they co-occurred with an enrichment of Tregs (*13*). Magen et al. identified a niche containing progenitor CD8^+^ T cells, mregDCs and CXCL13^+^ Th cells in ICI responders, while T cell rich non-responders were high in Tregs (*45*). CXCL13^+^ T cells were also associated with response to PD-1 blockade in an analysis by Sorin *et al.,* in contrast to DCs and Tregs that were found enriched in another community not significantly correlated to outcome (*56*). Yet, Chen *et al.* similarly identified immune hubs associated with response to PD-1 blockade that were rich in stem-like (progenitor exhausted) TCF7^+^ PD-1^+^ CD8^+^ T cell and mregDCs, also containing Tregs (*57*). This indicates that a careful definition of the Treg communities, including cell states (*21*) and local cytokine environments, will be required in order to identify accurately predictive biomarkers for resistance to PD-1/PDL1 blockade. Spatial analysis of pre-treatment biopsies from cohorts such as ARCTIC(*58*), MYSTIC(*59*), and CHECKMATE 227,(*60*) comparing anti-PDL1/PD-1 as monotherapy or in combination with anti-CTLA-4, could help to better define the Treg communities that confer resistance to blocking the PD-1/PD-L1 axis alone, yet may be responsive to the addition of Treg targeting.

Clinical trials for NSCLC that combine nivolumab or durvalumab anti-PD-(L)1 with ipilimumab or tremelimumab anti-CTLA-4 as mentioned above have shown no or only modest additional clinical benefit compared to monotherapy PD-(L)1 blockade (*61*). This may be due to lack of statistical power, by absence of a biomarker to select the patients that may benefit from additional CTLA-4 blockade. Interestingly, the MYSTIC trial showed that the combination of durvalumab and tremelimumab versus durvalumab monotherapy was performing worse across the whole patient population, but showed improved survival in a select patient group with high TMB(*22*). In line with this, we observed that there were two Treg communities with strong similarities to the Treg community we identified in mice, and which correlated with increased TMB, possibly representing communities that would be more responsive to CTLA-4 blockade or Treg depletion.

It should also be noted that it has been heavily debated whether the human anti-CTLA-4 antibodies ipilimumab and tremelimumab are Treg depleting. Treg depleting strategies are being explored clinically, such as an anti-CTLA-4 monoclonal antibody that was Fc-engineered to bind FcyRIIIA (botensilimab) (*62*), several different Treg depleting anti-CCR8 antibodies in Phase I/II studies, mogamulizumab, a CCR4 depleting antibody in Phase I/II studies (NCT02358473, NCT02946671) (*63*), and in preclinical studies like the Fc-optimised anti-CD25(*64*). One of the concerns with anti-CTLA-4 therapies is the enhanced toxicity seen in patients treated with ipilimumab or tremelimumab on top of anti-PD1 therapy. Local delivery of low-dose anti-CTLA-4 has been proposed to reduce toxicity while maintaining efficacy, such as recently shown for local delivery to the lymphatic basin in melanoma (*65*). However, the site for local delivery of a Treg depleting antibody therapy should be carefully considered. Like others before us (*26, 64*), we observed how the Treg depletion with anti-CTLA-4 was efficient in the tumors, but not occurring in the lymph nodes, most likely due to the differences in CTLA-4 expression. Of note, CTLA-4 levels are increased on activated Tregs (aTregs), which allows for selective depletion of these functionally suppressive Tregs (*65*), that are elevated both in tumors and tdLN. Intra- or peri-tumoral delivery might therefore be a more appropriate approach than systemic anti-CTLA-4 delivery (*44*).

In conclusion, we have used spatial cellular community analysis to investigate the nature of resistance to combined KRAS-G12C and PD-1 inhibition in a strongly immune evasive mouse model for NSCLC. We identified a cellular community that is rich in T cells and DCs, with evidence of local T cell activation and cytotoxicity, but that is inhibited by Treg-mediated suppression. Depletion of the Tregs led to profound reduction in tumor growth, longer survival and enhanced, and in some cases sustained, anti-tumor immunity. Analysis of treatment naïve patients showed that similar communities rich in CD4, CD8^+^ and Treg cells were found to co-occur with DCs in the peri-tumor space and correlated with increased TMB. We propose that a detailed spatial analysis of Treg rich communities in clinical samples from patients treated with KRAS-inhibitors, anti-PD-1/PDL1 or anti-CTLA-4 may provide the foundations for a very specific predictive biomarker.

## MATERIALS AND METHODS

### Study design

The goal of the study was to identify new targets within the spatial cellular communities in the tumour microenvironment of KRAS-G12C inhibitor treated mouse lung adenocarcinomas, to overcome therapy resistance. Previously published data from Imaging Mass Cytometry analysis of Lewis Lung Carcinoma treated for 7 days with KRAS-G12C inhibitors was reanalysed to look into the recurring cellular communities that could be potentially linked to therapy resistance. Relationships between cells within the communities were analysed, looking at cell-cell distances, expression of activation markers and neighbourhood enrichment. To validate the findings, a similar analysis was conducted in a human dataset of treatment naïve NSCLC tumour samples. The identified suppressor cells, namely Tregs, were then targeted in vivo using depleting antibodies, to verify that these were indeed instrumental in imposing resistance to KRAS-G12C + PD1 treatment.

### In vivo drug study and imaging mass cytometry

The IMC data was published previously in Van Maldegem, et al.(*9*) for dataset 1 and Mugarza, et al.(*8*) for dataset 2. For details on samples, staining, imaging and cell typing, please refer to the above references or Supplementary materials.

### Neighbour identification

As described in van Maldegem *et al* (*9*), cellular neighbours were identified during segmentation using the CellProfiler module ‘Measure Object Neighbours’, following steps to identify individual cells. Neighbours were identified if a cells boundary was within 15 μm (pixels) of the cell boundary of interest. Neighbour information was obtained for every cell object in each tissue.

### Identification of spatial cellular communities - mouse data

Each ‘neighbourhood’ was identified as a single cell and its local neighbours; defined by 15 μm object relationship output data from segmentation. For each cell, the proportion of each cell type (16 cell types for dataset 1 & 14 cell types for dataset 2) found in its neighbourhood was calculated (range: 0-1), as demonstrated by the equation below:

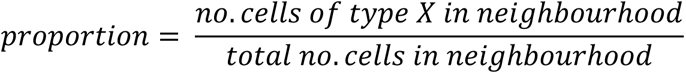

All cells per dataset were then clustered using Rphenograph, based on the neighbour proportion values calculated. A k=250 clustering input yielded 62 neighbourhood communities for dataset 1 and 254 communities for dataset 2.

Dimensionality reduction such as the R implementation of tSNE (RtSNE) (Van der Maaten *et al*, 2008), and dendrograms were used to determine which communities were similar and could therefore be analysed together for dataset 1. Agglomerative clustering using the AgglomerativeClustering function from the sklearn.cluster package in Python 3.9 was then used to group the 62 communities into 18. The clustree package was used to represent the merging of 62 communities into 30 and subsequently 18. Agglomerative clustering was also used to group 254 communities into 18, so that analysis of communities from datasets 1 and 2 were equal and thus the datasets could be analysed in parallel. For validation of the method to identify communities, please refer to Supplementary Materials.

### Pearson correlation

For comparing cell type relationships per ROI, the proportion of each cell type contributing to each ROI was calculated. For comparing cell type relationships within communities, the proportion of each cell type contributing to each of the 18 communities was calculated. For both instances this was followed by a Pearson correlation calculation of each cell type pair across all communities. A significant correlation was depicted by a p-value below 0.05, but significance was tiered as * = p < 0.05, ** = p < 0.01, *** = p < 0.001. Cell types were clustered based on their correlations.

### Neighbourhood enrichment analysis

As described in van Maldegem *et al*. (*9*), the neighbouRhood method developed by the Bodenmiller lab (https://github.com/BodenmillerGroup/neighbouRhood (*19*)) was used to identify the enrichment of cell types within the 15-pixel neighbourhood of each cell compared to random permutations of events, with a modification of only calculating enrichment in the CD8^+^ T cell neighbourhood, separated by community and using 1000 rounds of permutation. This was carried out for the community names T cell/normal adjacent community (T/NA), T cell/Type 1 macrophage community (T/M1), T cell/Dendritic cell community (T/DC), T cell/Type 2 macrophage community 1 (T/M2_1) and T cell/Type 2 macrophage community 2 (T/M2_2) communities for the MRTX1257 treatment setting. Neighbourhood enrichment analysis was also used for exploring cell pair relationships in the presence or absence of Tregs within the T/DC and T/M2_2 communities following MRTX1257 treatment. Enrichment scores were only deemed statistically significant if the p-value when comparing real neighbours to randomised neighbours was ≤ 0.01.

### Distance calculations

X- and Y-coordinates based on the centre of each cell, as generated through segmentation in CellProfiler, where used to calculate distances between cells. The cKDTree function from scipy.spatial package in Python 3.9 was used to compute distances of Dendritic cells and CD103^+^ Dendritic cells to CD4^+^, CD8^+^ and Tregs that were PD-1^+^ with a distance threshold of 800 pixels within the T/DC community following MRTX1257 treatment.

### Identification of spatial cellular communities – RUBICON data

The community identification method, developed by Schurch *et al.,* 2020, was applied to 141 non-small cell lung cancer tumour cores that were imaged with the Pan-Immune panel and 137 tumour cores that were imaged with the T cells & Stroma panel to identify groups of cells that commonly localised near one another.

Briefly, the method is as follows: A window was defined around every cell in an image and its 10 nearest neighbouring cells including the centre cell. These windows were clustered by their composition with respect to the 18 cell types in the Pan-Immune panel and the 20 cell types in the T cells & Stroma panel (with at least 10 cells on average per image) using MiniBatchKMeans. We optimised the parameters of the Schurch *et al.* method and identified 10 spatial cellular communities from the Pan-Immune panel. Similar to the spatial cellular communities reported in non-small cell lung cancer by Sorin *et al.,* 2023, we detected 30 spatial cellular communities using the T cells & Stroma panel. Communities were then assigned representative names based on the enrichment of cell densities within them.

Spatial community identities were mapped onto segmented cells and visualised using Cytomapper (Eling *et al.,* 2021), which were then validated by a pathologist’s assessment of serial H&E-stained tissue sections.

The cell density of spatial cellular communities was calculated by taking the number of cells assigned to a spatial cellular community divided by the total tissue area (cells/mm^2^).

### Association between cell densities of cell subtypes (T cells & Stroma panel) and spatial cellular communities (Pan-immune panel) – RUBICON data

We correlated the cell density of stroma-localised T cell subtypes detected using the T cells & Stroma antibody panel and the cell density of ten spatial communities detected using the Pan-immune antibody panel in cores from the same tumour region. For comparisons with multiple tumour cores/regions per tumour, we used linear mixed effects (LME) model analysis to incorporate patient ID as a random effect. We report the T-score and p-value of the model.

### Co-occurrence of spatial cellular communities – RUBICON data

The lowest quartile of the cell densities of Treg communities was approximately 25 cells/mm^2^. Therefore, we used 25 cells/mm^2^ as the threshold to determine if a spatial community was present or absent in a tumour core. We reported the proportion of paired tumour cores (T cells & Stroma panel, Pan-immune panel) with at least 25 cells/mm^2^ in *p2_C1: Tumour border* communities and in *p1* Treg communities.

### In vivo survival experiment

3LL-ΔNRAS(*38*) were cultured in RPMI medium supplemented with 10% fetal calf serum, 4 mM L-glutamine (Sigma-Aldrich), penicillin (100 U/ml), and streptomycin (100 mg/ml; Sigma-Aldrich). Cell lines were tested for mycoplasma and authenticated by short-tandem repeat DNA profiling by the Francis Crick Institute Cell Services Facility. Cells were allowed to grow for not more than 20 subculture passages.

Intravenous tail-vein injections of 10^6^ 3LL-ΔNRAS cells were carried out for orthotopic studies using 8-12 week old C57BL/6J mice. Mice were euthanised, with an overdose of pentobarbitone, when a humane endpoint of 15% weight loss was reached or any sign of distress was observed (i.e. hunched, piloerection, difficulty of breathing). In addition, if a mouse was observed to have a tumor burden in excess of 70% of lung volume when assessed by micro-CT scanning, they were deemed at risk of rapid deterioration in health and euthanised immediately.

Mice were anesthetized by isoflurane inhalation and scanned using the Quantum GX2 micro-CT imaging system (PerkinElmer) at a 50-μm isotropic pixel size. Serial lung images were reconstructed and analysed using Analyze12 (AnalyzeDirect) as previously described in Zaw *et al*, 2022.

BMS antibodies anti-PD-1 (clone 4H2, g1-D265A), and anti-CTLA-4 (clone 9D9, mlgG2a), with mlgG1-D265A and mlgG2a Isotope controls were given twice weekly at 200μg/dose by intraperitoneal injection. MRTX849 (adagrasib) was given by oral gavage daily at 100 mg/kg for a total of two weeks.

### Flow Cytometry

Flow cytometry was performed as previously (*8*) using the antibody mixes listed in Supplementary Table 1. Details of staining protocol, data acquisition and analysis can be found in Supplementary Materials.

## List of Supplementary Materials

Supplementary Materials text

Supplementary Table 1

Supplementary Figures 1-8

## Acknowledgements

We thank the science technology platforms at the Francis Crick Institute including Biological Resources, Scientific Computing, Bioinformatics and Biostatistics, and Flow Cytometry.

## Author contributions

Study design: MC, FvM, JD

Interpreting results: MC, FvM, JD

Writing manuscript: MC, FvM, JD

Acquisition and analysis of RUBICON human lung cancer dataset: CL, EC, MA, KSSE, AMa, CS

In vivo drug treatment experiments and analysis: PA, EM, SR, MM-A, CM

Performed and advised on data analysis: MC, FvM, MJ, KV, Amu

Statistical analysis: GK

Immunological expertise: TDdG

Manuscript revision and review: All

## Data availability

Dataset 1: https://hdl.handle.net/10779/crick.c.5270621.v2

Dataset 2: https://doi.org/10.25418/crick.19590259

## Code availability

- Link to Github page

## Supplementary Material

## SUPPLEMENTARY METHODS

### In vivo drug study and imaging mass cytometry

IMC data was published previously in van Maldegem, et al. (*9*) for dataset 1 and Mugarza, et al.(*8*) for dataset 2 In brief, 10^6^ ΔNRAS 3 Lewis lung carcinoma cells (*38*) were injected into the tail vein of 9-11 week old C57BL/6 mice. Following 3 weeks, mice were treated with either 50mg/kg MRTX1257 or Vehicle for 7 consecutive days. Mice were sacrificed on day 8 and lungs were harvested and frozen. Tumors of 3 mice from each treatment group for dataset 1 and 4 mice from each treatment group for dataset 2 were processed for imaging mass cytometry. Tissue slices were stained with a cocktail of antibodies conjugated to heavy metal isotopes and images were obtained using a Hyperion Imaging Mass Cytometer (Standard BioTools). Different antibody panels were used for the generation of datasets 1 and 2 (Supplementary Figure 1a, b). In total, six images were acquired from both Vehicle and MRTX1257 groups for dataset 1, and the Vehicle group for dataset 2, and five images were acquired from the MRTX1257 group for dataset 2. Further information regarding the in vivo study, tissue processing, antibody staining, and image acquisition can be found in van Maldegem, et al., 2021 (*9*).

### Image segmentation

Segmentation for the image sets from cohorts 1 and 2 was carried out using CellProfiler v3.1.9, including custom modules by Bodenmiller (https://github.com/BodenmillerGroup/ImcPluginsCP) and Ilastik v1.3.3b1. For cohort 1, a sequential segmentation pipeline was run to identify individual cells from the IMC images. In brief, probability maps were created in Ilastik for the separate identification of lymphocytes, macrophages, fibroblasts, tumor cells and endothelium, while remaining cell objects were identified using a nuclei expansion of 1-pixel. This pipeline also involved domain segmentation through generation of domain probability maps, enabling normal lung tissue, tumor tissue, and the interface region between normal and tumor sections to be identified. See van Maldegem *et al*.(*9*) for further details on the sequential segmentation strategy. For cohort 2, a segmentation pipeline was run which involved a 1-pixel expansion from the cellular nuclei to identify the cell objects. This method did not involve generation of domain information per tissue. For generation of both datasets, segmentation was run using imcyto (https://github.com/nf-core/imcyto). See ‘Data availability’ for details of CellProfiler modules used and project files generated for both segmentation methods.

### Normalisation, scaling, and clustering

Expression values for each marker were normalised to the mean intensity of Xenon134. Following this, the data for each image across both treatment groups was concatenated, creating a size of 282,837 cells for dataset 1 and 626,070 cells for dataset 2. Each channel was then scaled to the 99^th^ percentile. Mean pixel intensity of 17 cellular markers for dataset 1 (αSMA, B220, CD103, CD11c, CD3, CD44, CD45, CD4, CD68, CD8, EPCAM, F480, LY6G, MHCII, NKp46, PECAM, PVR) and 14 cellular markers for dataset 2 (αSMA, B220, CD103, CD11c, CD3, CD44, CD45, CD4, CD68, CD8, F480, Foxp3, MHC-II and NKp46) were used for clustering using Rphenograph (*66*) with k=20 to identify 30 clusters for dataset 1 and 33 clusters for dataset 2. These clusters formed the basis to identify cell types present in the tissue, including tumor, lymphocytes, myeloid cells, fibroblasts, and endothelium. More details available in van Maldegem *et al*. (*9*) for dataset 1 and Mugarza *et al*. (*8*) for dataset 2.

### Validation of method to identify communities

For validation using different input parameters, clustering was run on the neighbourhood proportion information for dataset 1 with a k-input value of 250 to produce 62 communities (as described in ‘Community detection’ section) and repeated for a k-input value of 350 to yield 47 communities. A tSNE was then used for dimensionality reduction of the cell type proportions contributing to each of the total 109 communities.

For validation across datasets, only the neighbourhood proportion values for the cell types that were shared across datasets 1 and 2 were used, comprising B cells, Dendritic cells, Dendritic cells CD103, Fibroblasts, Macrophages type 1, Macrophages type 2, NK cells, T cells CD4, T cells CD8, T reg cells and Tumor cells. Although both datasets contained cells labelled as Unclassified, they were not included in this analysis due to not being associated with a particular cell phenotype, rather lack of, and therefore could not be directly compared across the two datasets. These neighbourhood proportion values were clustered using Rphenograph with a k-input value of 250, for dataset 1 and 2 separately to yield 18 communities for both dataset 1 and dataset 2 after agglomeration. tSNE was then used for dimensionality reduction of the cell type proportions contributing to each of the total 36 communities.

### Flow cytometry

Mouse lung tumors were cut finely and incubated with digestion solution (collagenase 1 mg/ml; ThermoFisher and DNase I 50 U/ml; Life Technologies) at 37°C for 45 minutes. Cells were then filtered through 70 μm strainers (Falcon) and red blood cells were lysed using ACK buffer (Life Technologies). After washes in PBS, cells were stained with fixable viability dye eFluor870 (BD Horizon) for 30 minutes at 4°C and blocked with CD16/32 antibody (BioLegend) for 10 minutes. Samples were then washed three times in FACS buffer (2 mM EDTA and 0.5% bovine serum albumin in PBS, pH 7.2) before staining of surface markers using fluorescently labelled antibody mixes (See Supplementary Table 1). Cells were fixed with a Fix/lyse solution (eBioscience) after staining. If intracellular staining was carried out, cells were instead fixed with Fix/Perm solution (Invitrogen), followed by intracellular antibody staining. Samples were then resuspended in FACS buffer and analysed using a FACSymphony analyser (BD). Data was analysed using FlowJo.

**Supplementary Figure 1.**
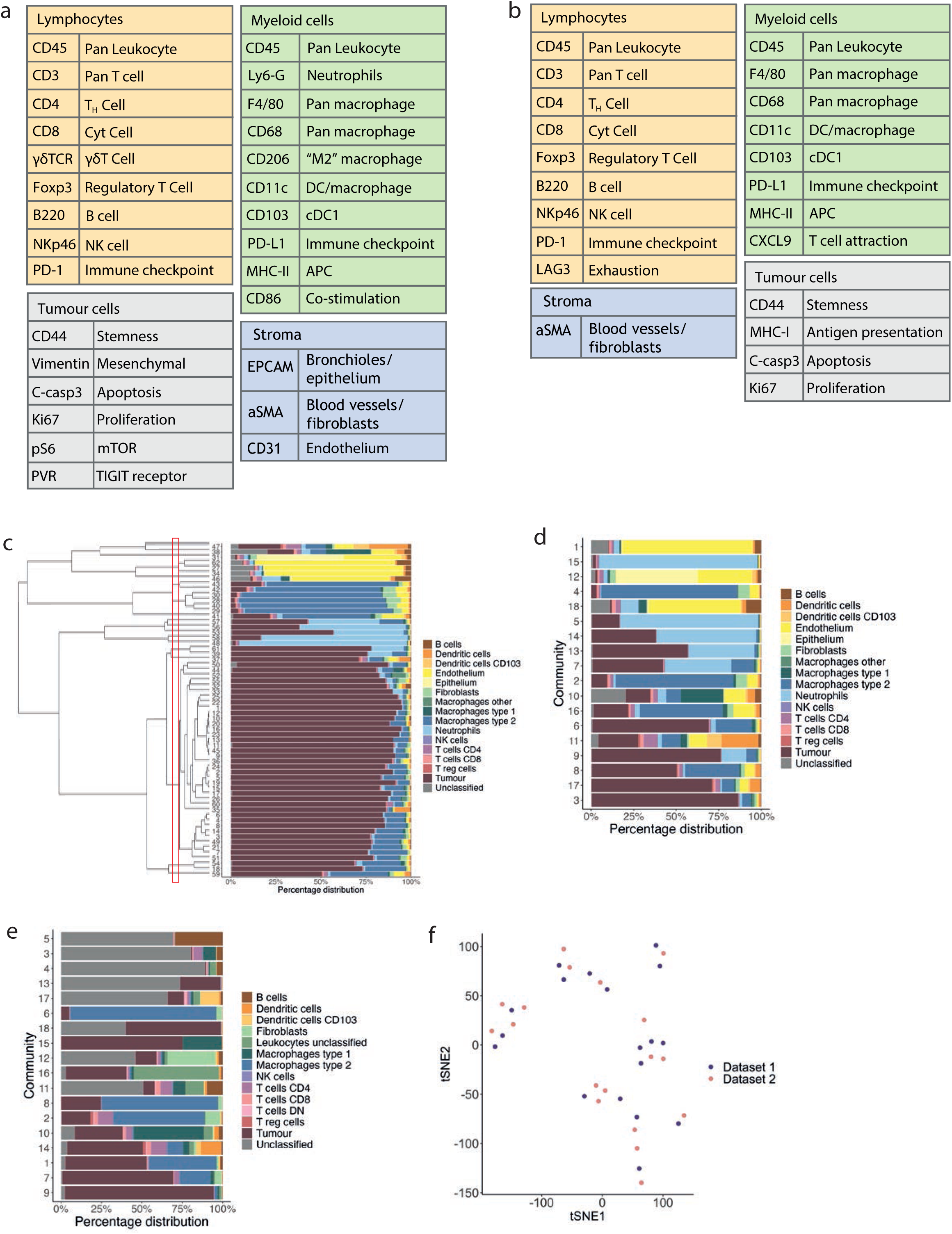
a) Panel of a) 27 antibodies for dataset 1 and b) 21 antibodies for dataset 2 that identify multiple cell types across lymphocyte, myeloid, tumour and stromal compartments, as well as immune checkpoint markers and cell phenotypic markers including for detection of proliferation and apoptosis. Additional antibodies for the dataset 2 panel (b) include LAG-3 for identifying exhausted T cells, MHC-I and CXCL9, a chemokine associated with T cell attraction. c) Percentage distribution of cell types contributing to 62 spatial communities identified from Rphenograph clustering of neighbourhood information with k=250. Dendrogram of how 62 communities were agglomerated to 18 communities, with the 18 community point boxed in red. d) Percentage distribution of all cell types assigned to each of the 18 communities following clustering of neighbourhood information for dataset 1. e) Percentage distribution of all cell types assigned to each of the 18 communities following clustering of neighbourhood information for dataset 2. f) tSNE plot of the 18 communities generated for dataset 1 and 18 communities generated for dataset 2 following Rphenograph clustering based on the proportion of only the cell types shared across both datasets within the neighbourhood of each cell.

**Supplementary Figure 2.**
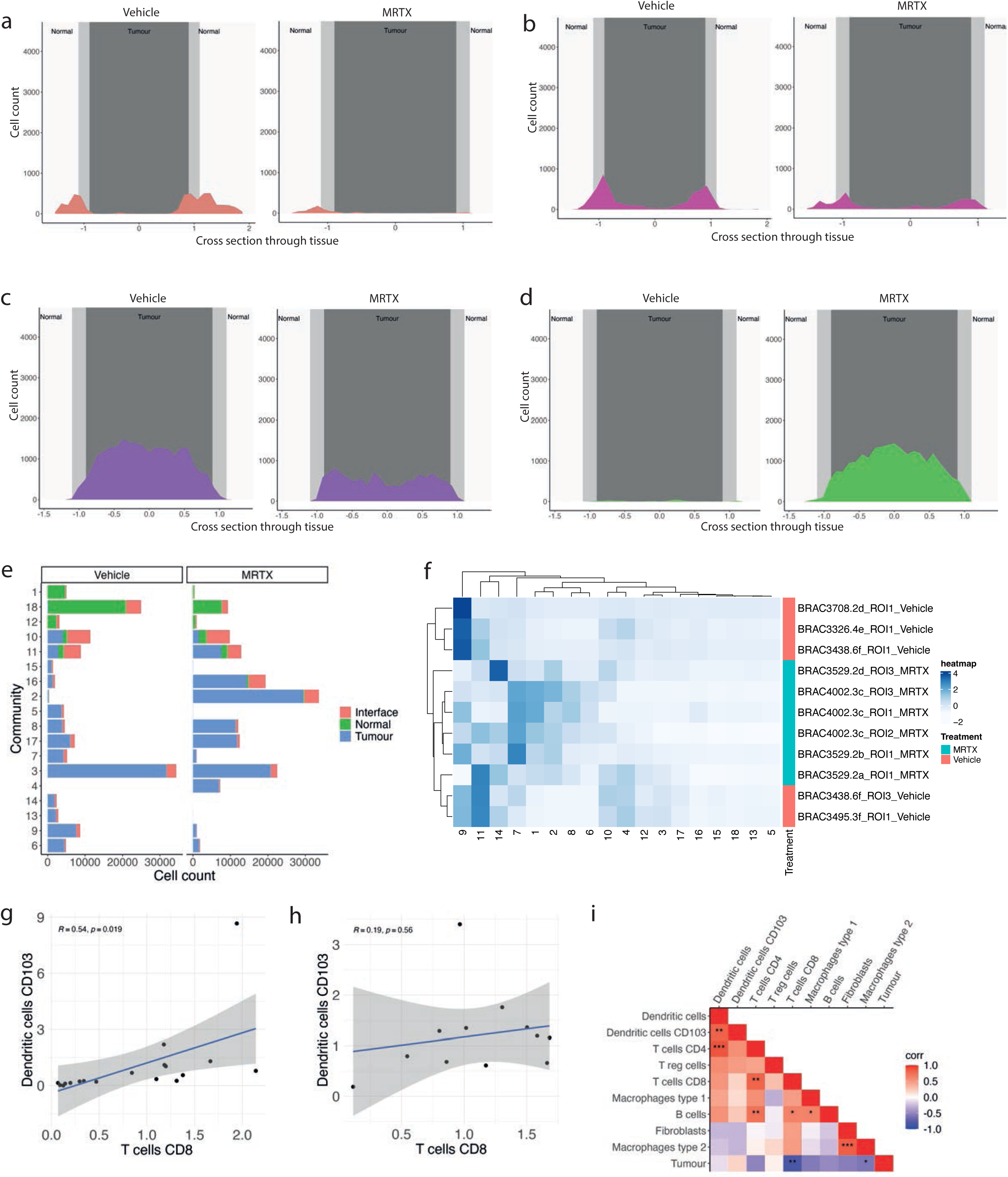
a) Cell count of community relative to cross section through the tissue, where 0 represents the centre point of the tumour for community a) 18 b) 10, c) 2 and d) 3, for Vehicle (left) and MRTX1257 (right) treatment settings. e) Percentage distribution of all 18 communities across normal, interface and tumour domains of the tissue for dataset 1. f) Hierarchical clustering of community proportion per ROI for dataset 2, with use of dendrogram to show relationships between similar ROIs, similar communities, and community distribution across the treatment groups. g) Pearson correlation calculation of the proportion of Dendritic cells CD103 and T cells CD8^+^in each of the 18 communities for dataset 1. h) Pearson correlation calculation of the proportion of Dendritic cells CD103 and T cells CD8^+^in each of the 12 ROIs for dataset 1. i) Pearson correlation calculation on the proportion of each cell type pair within each ROI. * = p < 0.05, ** = p < 0.01, *** = p < 0.001. Cell types clustered based on correlation value.

**Supplementary Figure 3.**
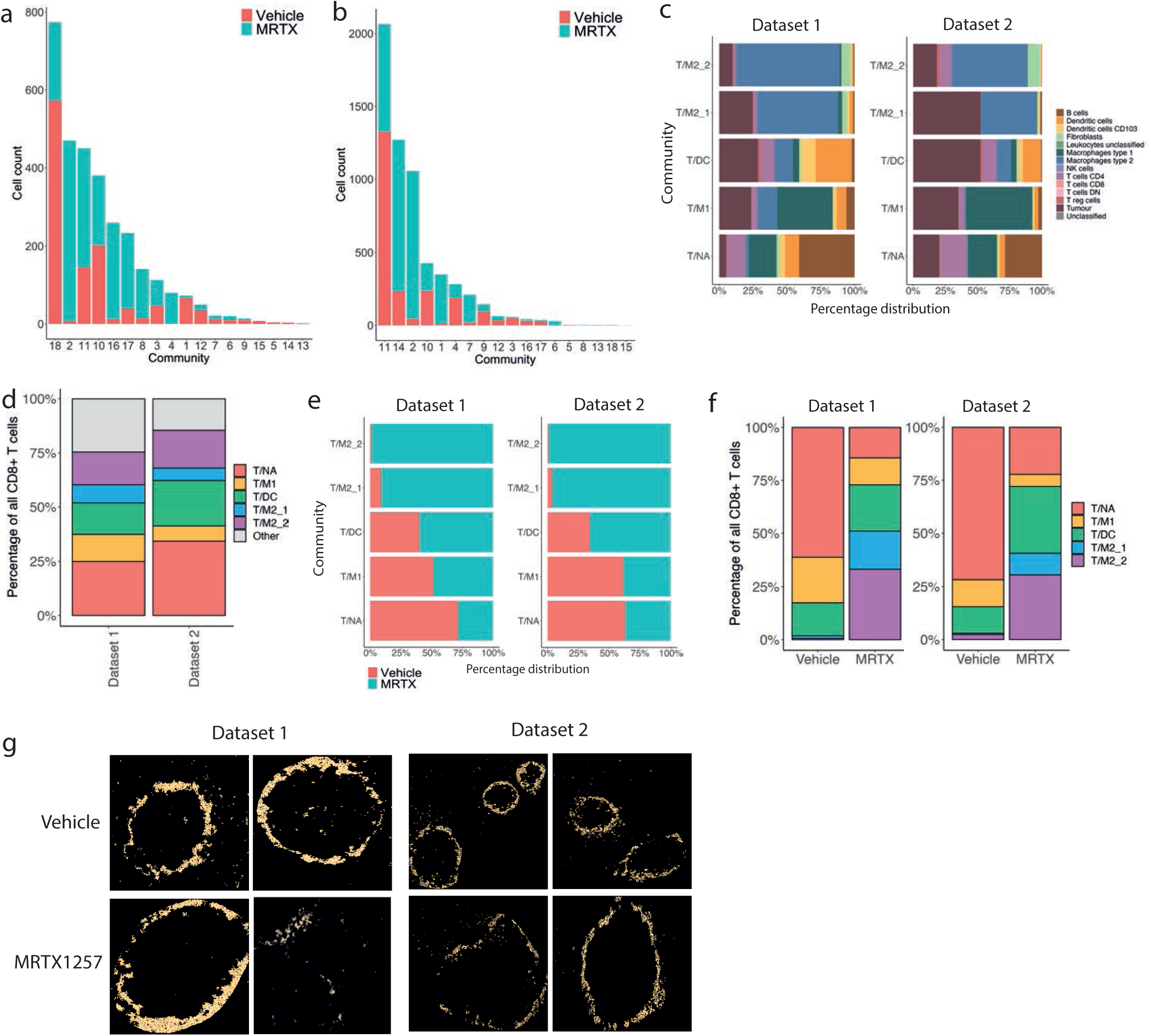
a) Number of CD8^+^ T cells assigned to each community, with bars coloured by distribution of those cells across Vehicle and MRTX1257 treatment groups for a) dataset 1 and b) dataset 2. c) Percentage distribution of cell types shared across datasets 1 and 2 contributing to T/NA, T/M1, T/DC, T/M2_1 and T/M2_2 communities. d) Percentage distribution of all CD8^+^ T cells in dataset 1 and 2 across T/NA, T/M1, T/DC, T/M2_1 and T/M2_2 communities and all ‘other’ communities. e) Distribution of all cells assigned to T/NA, T/M1, T/DC, T/M2_1 and T/M2_2 communities across Vehicle and MRTX1257 treatment groups, relative to the proportion of each treatment group across the whole cohort size for dataset 1 (left) and dataset 2 (right). f) Percentage of all CD8^+^ T cells found in the top 5 communities, coloured by their distribution across each of the top 5 communities in Vehicle and MRTX1257 treatment groups, for dataset 1 (left) and dataset 2 (right). g) Visualisation of cell outlines for cells assigned to community T/M1, from Vehicle and MRTX1257 treatment groups of datasets 1 (left) and 2 (right).

**Supplementary Figure 4.**
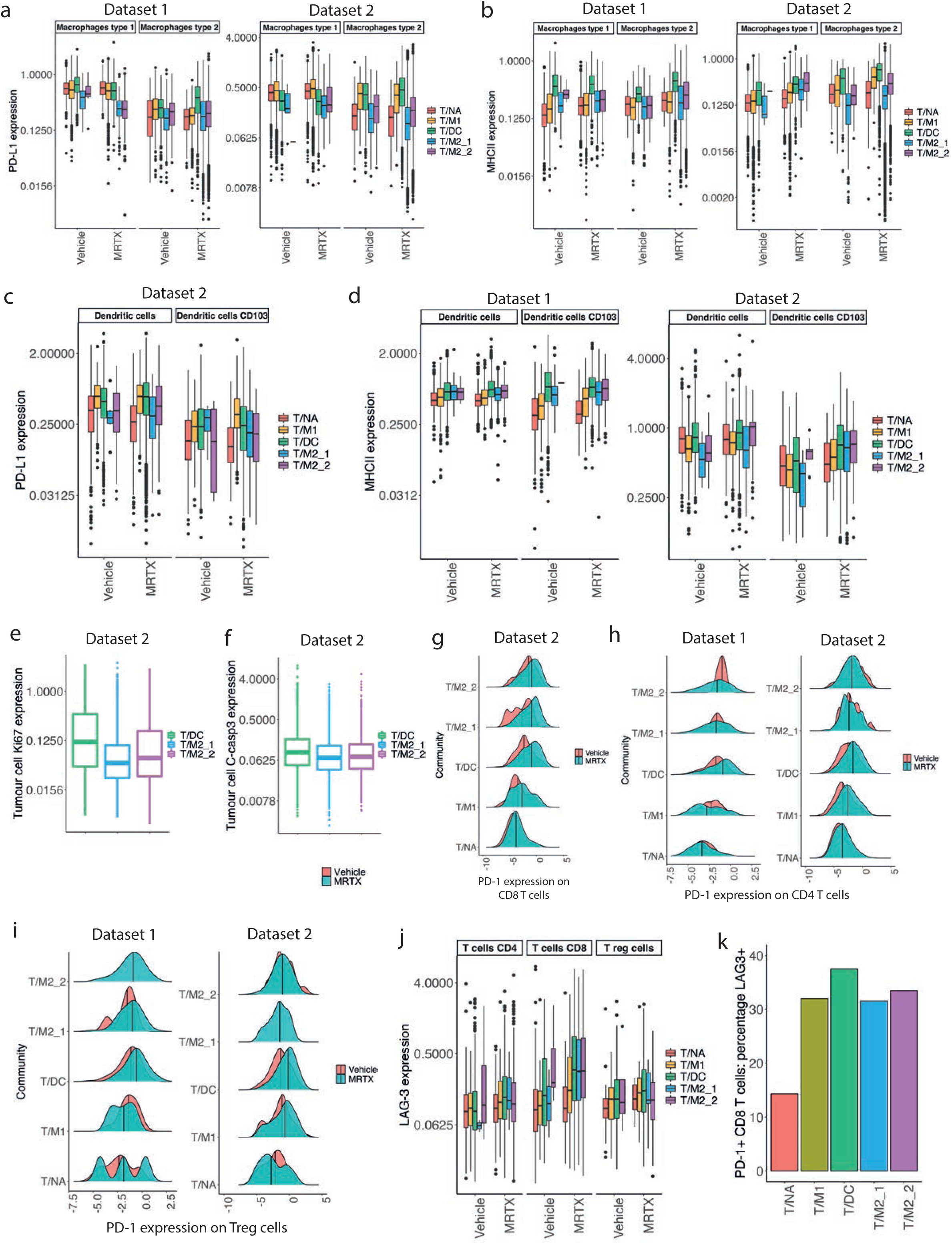
a) Mean expression of a) PD-L1 and b) MHC-II on macrophages type 1 and type 2 for dataset 1 (left) and dataset 2 (right) in T/NA, T/M1, T/DC, T/M2_1 and T/M2_2 communities for Vehicle and MRTX1257 treatment groups, values were log2 scaled. c) Mean expression of PD-L1 on dendritic cells and dendritic cells CD103 in communities A-E for Vehicle and MRTX1257 treatment groups, values were log2 scaled. d) Mean expression of MHC-II on dendritic cells and dendritic cells CD103 for dataset 1 (left) and dataset 2 (right) in T/NA, T/M1, T/DC, T/M2_1 and T/M2_2 communities for Vehicle and MRTX1257 treatment groups, values were log2 scaled. e) Mean expression of e) Ki67 and f) cleaved-caspase 3 (c-casp3) on tumour cells in communities C, D and E following treatment with MRTX1257 for dataset 2, values were log2 scaled. g) Mean expression of PD-1 on g) CD8^+^ T cells for dataset 2, h) CD4^+^ T cells for dataset 1 (left) and dataset 2 (right) and i) regulatory T cells for dataset 1 (left) and dataset 2 (right) in T/NA, T/M1, T/DC, T/M2_1 and T/M2_2 communities for Vehicle and MRTX1257-treated groups. j) Mean expression of LAG-3 on T cells CD4^+^,T cells CD8^+^and T reg cells in communities A-E for Vehicle and MRTX1257 treatment groups, values were log2 scaled. k) Percentage of PD-1^+^ CD8^+^ T cells that are positive for LAG-3 expression (based on a mean expression threshold of 0.5), across T/NA, T/M1, T/DC, T/M2_1 and T/M2_2 communities following MRTX1257treatment.

**Supplementary Figure 5.**
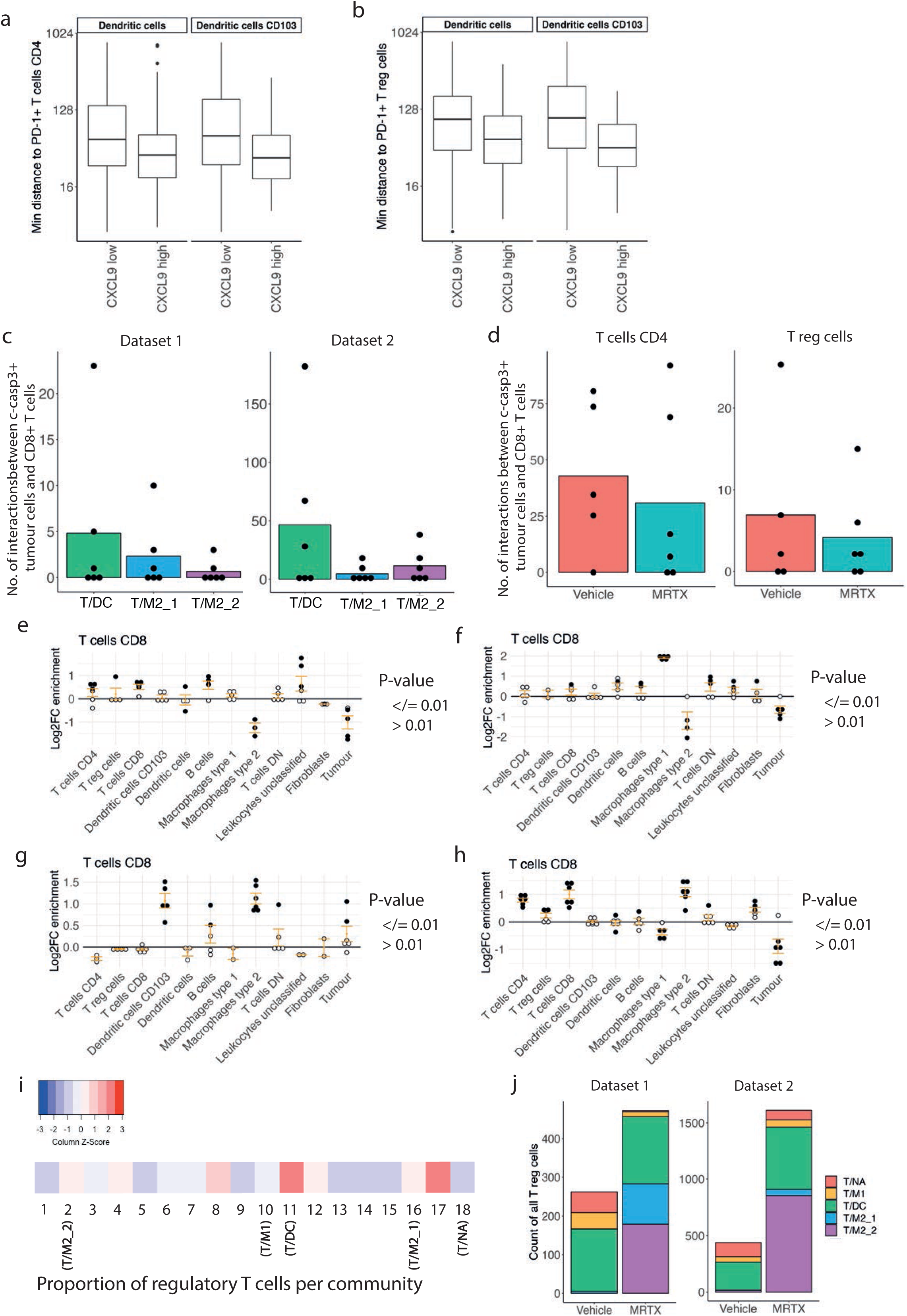
a) Minimum distance of dendritic cells and CD103^+^ dendritic cells that have ‘low’ or ‘high’ CXCL9 expression (based on a threshold of 0.5) to PD-1^+^ b) CD4^+^ T cells and c) regulatory T cells, within 800 pixels in the T/DC community, distance values were log2 scaled. c) Number of times a c-casp3^+^ tumour cell is found in the 15-pixel neighbourhood of a CD8^+^ T cells within T/DC, T/M2_1 or T/M2_2 communities following MRTX1257 treatment for dataset 1 (left) and dataset 2 (right). Values averaged across ROIs. d) Number of times a c-casp3^+^ tumour cell is found in the 15-pixel neighbourhood of a CD4^+^ T cell (left) and regulatory T cell (right), within the T/DC community, compared across Vehicle and MRTX1257 treatment groups for dataset 2. Count is relative to the proportion of tumour cells that were c-casp3^+^ in Vehicle vs MRTX1257 treatment groups and averaged across ROIs. e) Log2 fold changes in enrichment from neighbouRhood analysis for CD8^+^ T cells in e) T/NA, f) T/M1, g) T/M2_1 and h) T/M2_2 communities, following treatment with MRTX1257. Filled circles represent images from which enrichment value was statistically significant compared to randomisation of the spatial arrangements within all top 5 communities following treatment with MRTX1257. i) Proportion of regulatory T cells contributing to each of the 18 original communities for dataset 1. j) Count of regulatory T cells in the top 5 communities, split by Vehicle and MRTX1257 treatment groups for dataset 1 (left) and dataset 2 (right).

**Supplementary Figure 6.**
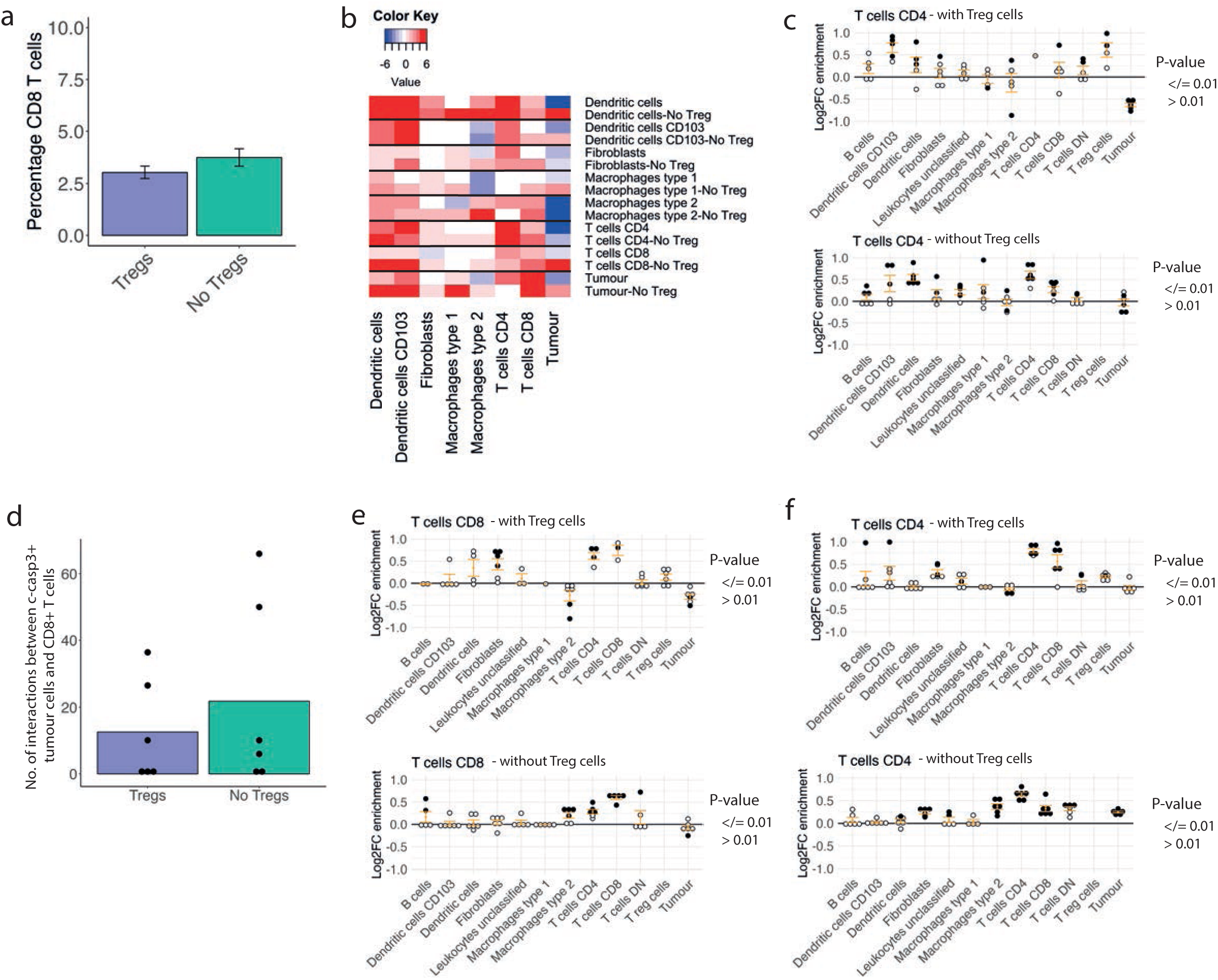
a) Percentage of CD8^+^ T cells within the ‘Tregs’ and ‘No Tregs’ neighbourhoods of T/DC community in MRTX1257 treatment group, averaged across ROIs. b) Heatmap of the enrichment scores from neighbouRhood enrichment analysis of each cell type in the neighbourhood of every other cell type in the ‘Tregs’ or ‘No Tregs’ neighbourhood for T/DC community in MRTX1257 treated samples. Score calculated individually per ROI and summed together. c) Log2 fold changes in enrichment from neighbouRhood analysis for CD8^+^ T cells in ‘Tregs’ (top) and ‘No Tregs’ (bottom) neighbourhoods within T/DC community following treatment with MRTX1257. Filled circles represent images from which enrichment value was statistically significant compared to randomisation of the spatial arrangements within T/DC community following treatment with MRTX1257 for dataset 2. d) Number of times a c-casp3^+^ tumour cell is found in the 15-pixel neighbourhood of a CD4^+^ T cell within T/DC community, compared across ‘Tregs’ and ‘No Tregs’ neighbourhoods in dataset 2, averaged per ROI following MRTX1257 treatment. Count is relative to the proportion of tumour cells that were c-casp3^+^ in ‘Treg’ vs ‘No Treg’ groups. e) Log2 fold changes in enrichment from neighbouRhood analysis for **e)** CD8^+^ T cells and **f)** CD4^+^ T cells in ‘Tregs’ (top) and ‘No Tregs’ (bottom) neighbourhoods within T/M2_2 community following treatment with MRTX1257. Filled circles represent images from which enrichment value was statistically significant compared to randomisation of the spatial arrangements within the T/M2_2 community following treatment with MRTX1257 for dataset 2.

**Supplementary Figure 7.**
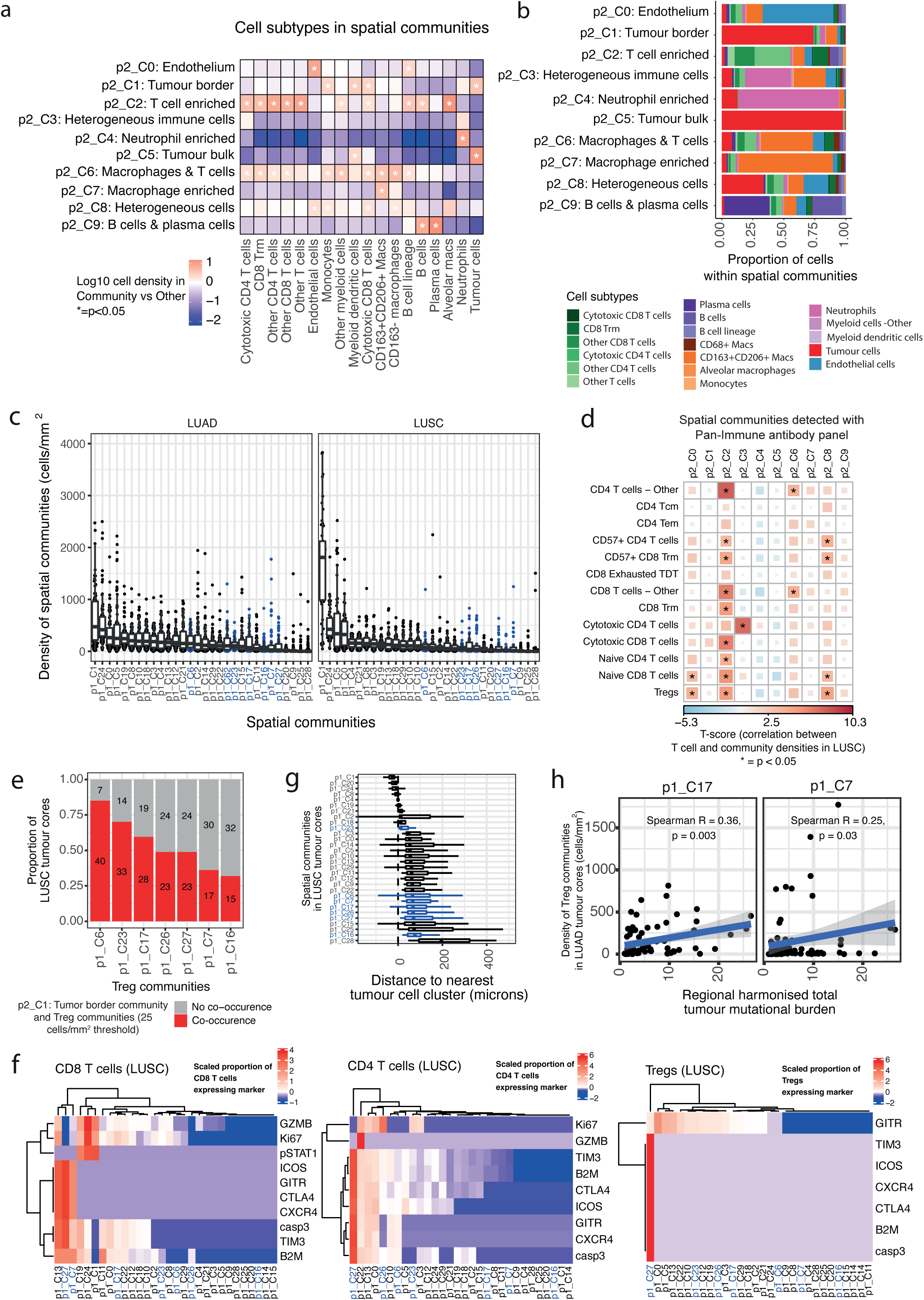
a) Ten spatial communities detected in 141 tumour cores from 69 patients with non-small cell lung cancer. The log10 cell density of cell subtypes in one community compared to all other communities is displayed. Linear mixed effects model (LMEM) to test the association between the density of cell subtypes with one community compared to that of other communities, with patient as a random covariate to adjust for multiple cores per tumour. Analysis of variance test comparing the LMEM to the null model, p-values are unadjusted. *:p < 0.05 (*20*). b) Proportion of cell subtypes assigned to each of the ten spatial communities. 141 tumour cores from 69 patients. c) Density of spatial cellular communities detected in LUAD and LUSC tumour cores. 70 LUAD cores from 41 patients, 49 LUSC cores from 21 patients. d) Correlation between the density of stroma-localised T cell subtypes detected using the T cells & Stroma antibody panel and the cell density of ten spatial communities detected using the Pan-immune antibody panel, in 47 LUSC tumour cores from 22 patients. LMEM to test the association between T cell density and cell density of communities with patient as a random covariate to adjust for multiple cores per tumour. Analysis of variance test comparing the LMEM to the null model, p-values are unadjusted. *:p < 0.05. e) Proportion of LUSC tumour cores that contain at least 25 cells/mm^2^ of Treg communities (*p1_C6, p1_C7, p1_C16, p1_C17, p1_C23, p1_C27*) *and p2_C1: Tumour border* communities. 47 LUAD tumour cores from 22 patients. f) Heatmap displaying the scaled proportion of CD8^+^ T cells, CD4^+^ T cells and Treg cells expressing phenotypes of interest. 51 LUSC tumour cores from 23 patients. g) Per-image median distance between cells of a community and their nearest tumour cell cluster. Tumour cell clustering method described in Magness *et al.,* manuscript in review. 51 LUSC tumour cores from 23 patients. h) Spearman correlation between the density of Treg communities and total harmonised tumour mutational burden. 70 LUAD cores from 41 patients. LUAD, lung invasive adenocarcinoma; LUSC, squamous cell carcinoma. Treg communities are highlighted in blue.

**Supplementary Figure 8.**
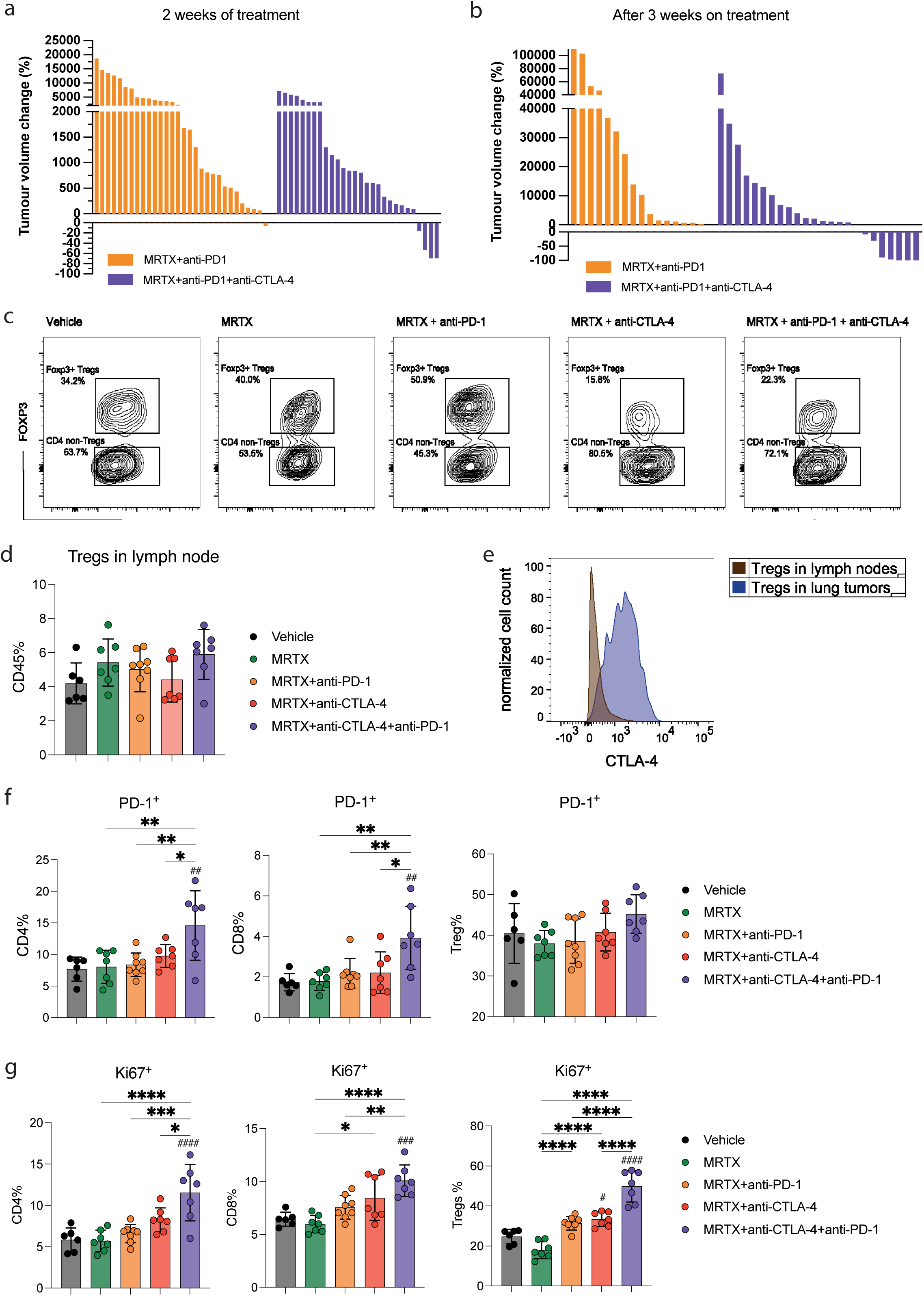
a) Tumour volume changes after **a)** two weeks and **b)** three weeks of treatment as measured by μCT scanning, for MRTX1257+anti-PD-1 and MRTX1257+anti-PD-1+anti-CTLA-4 treatment groups. c) Representative flow cytometry contour plots for FOXP3^+^ CD4^+^ Treg cells in each treatment group. d) Percentage of all CD45^+^ cells identified as regulatory T cells (gated as CD45^+^ CD3^+^ CD4^+^ Foxp3^+^) measured by flow cytometry in the tumour draining lymph nodes (each dot represents a mouse, one-way ANOVA). e) Histogram of CTLA-4 expression on FOXP3^+^ Tregs following MRTX1257treatment in the tumour (blue) or tumour draining lymph node (brown). f) Percentage of CD4^+^ T cells, CD8^+^ T cells and Treg cells that are PD-1^+^, measured by flow cytometry in the lymph nodes. g) Percentage of CD4^+^ T cells, CD8^+^ T cells and Treg cells that are Ki67^+^, measured by flow cytometry in the lymph nodes.

**Supplementary Table 1.**
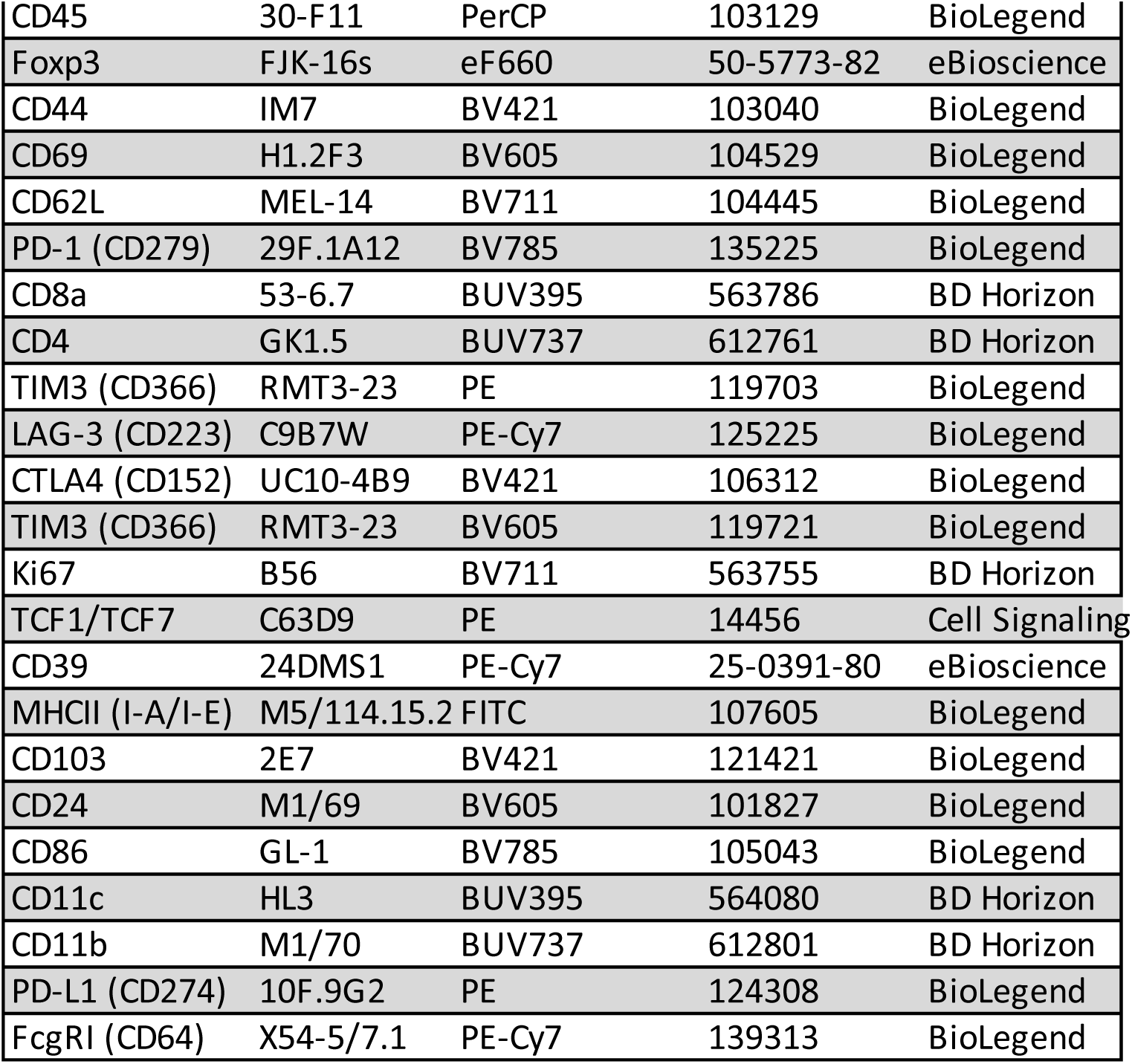
Antibodies used for flow cytometry.

